# Alternative splicing switches human Musashi-2 between cytoplasmic and nuclear regulatory functions

**DOI:** 10.64898/2026.05.16.725683

**Authors:** Kathryn Walters, Nadine Koertel, Amber Baldwin, Rachel Todebush, Marcin P. Sajek, Neelanjan Mukherjee

## Abstract

Musashi-2 (MSI2) is an RNA-binding protein that controls self-renewal and differentiation in the hematopoietic and nervous systems and sustains myeloid leukemia. It both activates and represses translation of its target mRNAs, outcomes usually attributed to cellular context. We find instead that the capacity to activate translation is determined by an alternative 3’ splice acceptor in exon 12, which splits full-length MSI2 into isoforms that carry or lack an 18-amino-acid insertion in the C-terminal disordered region. MSI2-328 is cytoplasmic and the insertion-containing MSI2-324 is nuclear. The insertion is necessary and sufficient for nuclear localization. Tethering MSI2-328 to a reporter mRNA promotes translation, whereas tethering MSI2-324 does not, and removing the insertion restores activation. Consistent with their localization, MSI2-328 binds translation factors and 3’ UTRs, while MSI2-324 binds splicing factors and introns. Use of the insertion-generating acceptor ranges from 19% in skeletal muscle to 51% in the brain and increases in several tumor types. Altogether, one splicing event toggles MSI2 between the cytoplasm and the nucleus, so localization, rather than cellular context, determines its regulatory function.

## INTRODUCTION

The Musashi family of RNA-binding proteins, consisting of MSI1 and MSI2, regulate stem cell maintenance, neural development, and cancer progression ^1,2^. These proteins share 75% sequence similarity ^3^. They contain two highly conserved RNA recognition motifs (RRMs) that bind UAG-containing elements within target mRNAs and have overlapping RNA-binding specificity and partial functional redundancy ^3–8^. Despite these similarities, MSI proteins have been reported to exert diverse and sometimes opposing effects on gene expression.

MSI1 was initially identified as a translational repressor of *Numb* mRNA ^9,10^, but subsequent studies have shown that it can both enhance and repress translation of its targets, sometimes within the same cellular context ^6,11–13^. Likewise, MSI2 has been reported to enhance mRNA stability ^10,14–17^, to promote translation without altering stability ^1,18,19^, to promote decay without affecting translational efficiency ^20^, and to repress translation ^21,22^. Two mechanisms have been proposed for Musashi-mediated translational regulation. Competition with eIF4G for PABPC1, which represses translation, was established for MSI1 ^23^, and activation through the poly(A) polymerase GLD2 was established in Xenopus ^24^. However, neither mechanism has been demonstrated for human MSI2. These diverse regulatory activities have instead been described as context-specific, without identifying what distinguishes one context from another ^1,19^.

Although Musashi proteins have been characterized primarily as regulators of mRNA translation and stability, they have also been implicated in nuclear RNA processing. Musashi binding downstream of alternative exons promotes exon inclusion, and combined loss of MSI1 and MSI2 results in widespread changes in alternative splicing ^25,26^. Musashi proteins therefore act in both compartments, and their differential localization is a possible explanation for their diverse regulatory activities. Neither what determines MSI2 localization nor how it shapes regulatory activity has been examined.

Alternative splicing generates RNA-binding protein isoforms with distinct regulatory functions, including differences in subcellular localization ^27,28^. MSI2 is alternatively spliced ^3,29^ and its isoforms are not functionally equivalent in triple-negative breast cancer ^30^. Nevertheless, nearly all functional work on human MSI2 has used a single 328 amino acid isoform (ENST00000284073). Whether alternative splicing controls MSI2 subcellular localization and contributes to its context-dependent regulatory activities remains unknown.

Here, we show that alternative splicing regulates MSI2 function by controlling its subcellular localization. An alternative 3′ splice acceptor in exon 12 introduces an 18-amino-acid insertion that directs the resulting isoform to the nucleus, whereas the isoform lacking the insertion localizes predominantly to the cytoplasm and promotes translation. Removing the insertion restores cytoplasmic localization and translational activation. Consistent with their localization, the two isoforms associate with different cofactors and bind different transcript regions. The nuclear isoform preferentially binds introns, in line with the splicing function attributed to Musashi proteins ^25,26^. Use of the acceptor that generates nuclear MSI2 ranges from 19% in skeletal muscle to 51% in brain and increases in several tumor types, so the balance between the isoforms varies with biological context. Altogether, these findings provide a molecular basis for the context-dependent regulatory activities attributed to MSI2.

## RESULTS

### An alternative exon 12 acceptor distinguishes the two endogenous MSI2 proteins

To investigate the contribution of alternative splicing on MSI2 function, we first examined endogenous MSI2 protein expression via western blotting. Using a panel of cell lines used to characterize MSI2 and reported to differ in its regulatory activity ^6,18,21,23,31–36^, we resolved two MSI2 species approximately 2 kDa apart in all cell lines tested (Fig 1A). Similar patterns of two MSI2 protein species have been reported previously ^18,33^. These observations prompted us to examine which alternatively spliced transcripts might account for the two predominant protein species.

**Figure 1.**
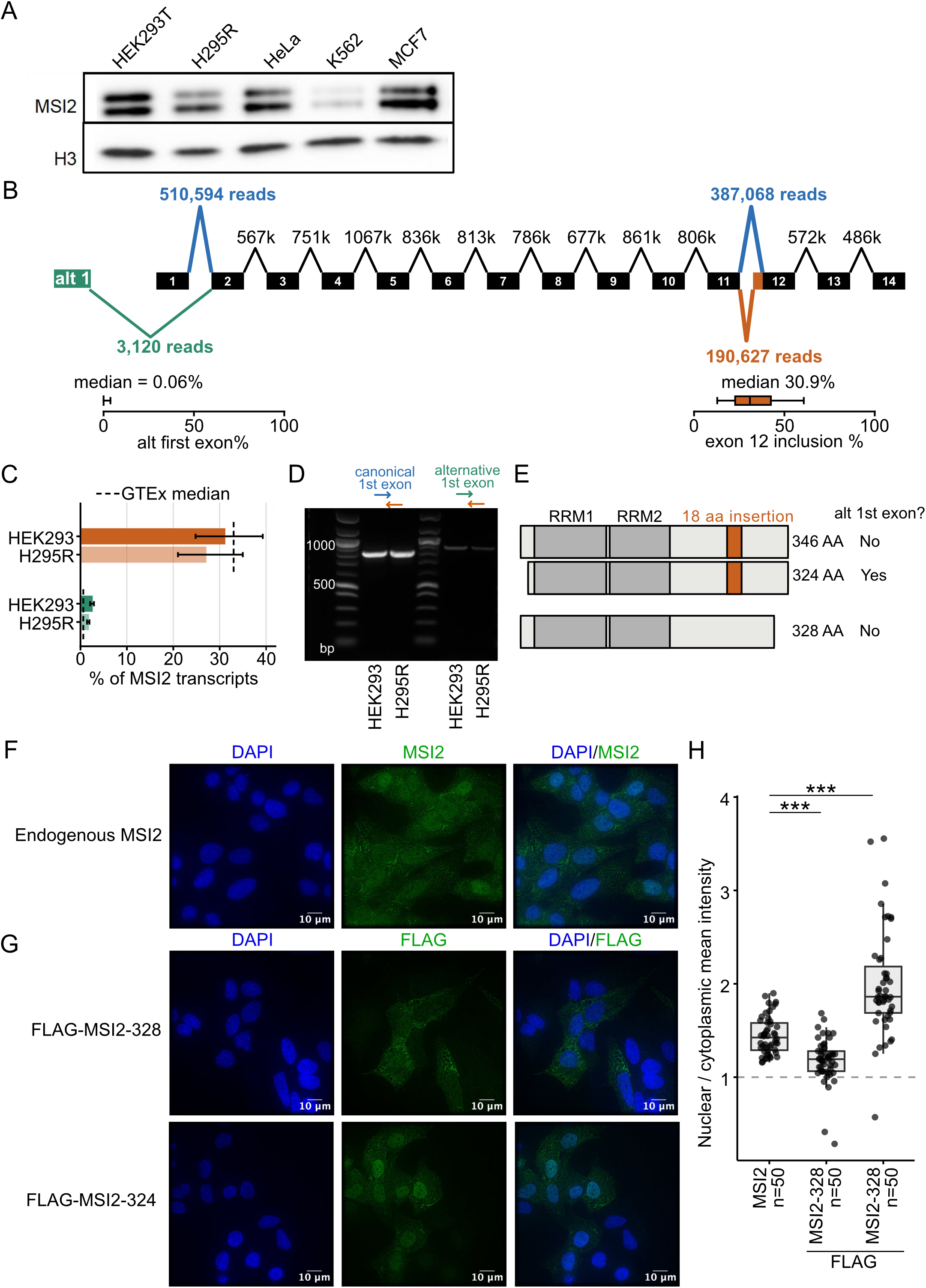
Alternative splice site selection generates distinct MSI2 isoforms with different subcellular localization. (A) Western blot of endogenous MSI2 across the indicated human cell lines. Histone H3 is used as a loading control. (B) Sashimi plot of MSI2 exons. Exons are shown as black boxes, the alternative first exon (green), the exon 12 insertion that adds 18 amino acids (orange), and the respective canonical splice-junctions for those events (blue). Splice-junction read counts and median alternative splice site usage across GTEx samples are indicated. (C) RT-qPCR quantification of MSI2 isoform composition in HEK293 and H295R cells. Bars show the fraction of full-length MSI2 transcripts that contains the exon 12 insertion and the alternative first exon, each as a percentage; the dashed line indicates the corresponding GTEx value. (D) Agarose gel with RT-PCR products from HEK293FT and H295R cells. Products were amplified using a reverse primer to the 18aa insertion, and forward primer located in either the canonical first exon (828 bp product) or alternative first exon (888 bp product). (E) Schematic of MSI2 protein isoforms arising from alternative first exon usage and exon 12 splice acceptor selection. RNA recognition motifs (RRM1 and RRM2), the 18-amino-acid insertion, and predicted protein lengths are indicated. (F) Representative immunofluorescence of endogenous MSI2 (green) and DAPI (blue). Scale bar, 10 μm. (G) Representative immunofluorescence of FLAG-MSI2-328 and FLAG-MSI2-324 (green) and DAPI (blue). Scale bar, 10 μm. (H) Nuclear-to-cytoplasmic ratio of mean intensity for endogenous MSI2 and dox-induced MSI2-328-FLAG and MSI2-324-FLAG. n indicates the number of cells imaged. Dashed line indicates a ratio of 1. Significance was tested by Mann-Whitney U test ***p < 0.001.

GENCODE annotates two full-length, protein-coding MSI2 transcripts, ENST00000284073 and ENST00000416426, which differ at the first exon and the 3’ splice acceptor of exon 12 (Sup Fig 1A). These transcripts encode proteins of 328 and 324 residues, which differ both at their N-terminus and by the presence of an 18aa sequence within the C-terminal intrinsically disordered region (IDR). A third annotated protein-coding transcript, ENST00000322684, uses an alternative terminal exon within the canonical intron 10, producing a truncated 251-residue protein that terminates upstream of the 18-amino-acid sequence and is therefore not examined here (Sup Fig 1A).

To determine the relative usage of these alternative transcript features, we analyzed splice-junction reads across 19,788 GTEx RNA-seq samples. The alternative first exon of ENST00000416426 accounted for only 0.6% of junction reads at that position (Fig 1B and Sup Fig 1B). In contrast, exon 12 acceptor usage was more evenly distributed, with the upstream acceptor, which adds the 18 amino acids, used in 33% of reads and the acceptor 54 nucleotides downstream in 67% of reads (Fig 1B and Sup Fig 1B). We tested these observations in HEK293 and H295R cells by designing qPCR primers to distinguish the alternative first exons and exon 12 splice acceptors independently. Transcripts containing the alternative first exon represented only 2.5% and 1.7% of MSI2 transcripts. Transcripts including the exon 12 insertion accounted for 31.2% and 27.1% of full-length MSI2 mRNA, consistent with the frequencies observed in GTEx (Fig 1C).

Although the qPCR and splice-junction analysis independently quantify these two alternative splicing events, they do not establish whether the events occur on the same transcript. We therefore performed RT-PCR using primers spanning either the alternative or canonical first exon and the insertion-containing exon 12 splice acceptor. We detected products corresponding to both isoforms, with a stronger product observed for the canonical first exon (Fig. 1D). Together with the splice-junction and qPCR analyses, these findings support the presence of an insertion-containing transcript that predominantly uses the canonical first exon. This transcript is predicted to encode a 346-residue protein that differs from the canonical 328-residue protein only by the 18-amino-acid insertion, whereas the annotated 324-residue protein also lacks 22 N-terminal residues (Fig. 1E). We refer to these proteins as MSI2-328, MSI2-346, and MSI2-324, respectively. Although MSI2-346 is not currently among the annotated human transcripts, its sequence corresponds to the canonical 346-residue mouse Msi2 protein (UniProt Q920Q6-1) ^29^ and previous studies have also identified two mouse Msi2 species of approximately 35- and 37 kDa ^3,29^. The predicted mass difference between MSI2-328 and MSI2-346 is 1.77 kDa, consistent with the ∼2 kDa separation we observe in human cell lines and with that reported for the mouse species.

### MSI2 isoforms localize to different compartments

MSI proteins have both nuclear and cytoplasmic regulatory functions, so we asked whether the two MSI2 isoforms differ in localization. Immunofluorescence for endogenous MSI2, using an antibody against the C terminus that detects both isoforms, showed signal in both the cytoplasm and the nucleus (Fig 1F). A mixed localization pattern could arise from one protein distributed across both compartments or two proteins each largely confined to one. To distinguish these, we expressed FLAG-MSI2-328 and FLAG-MSI2-324 individually. We used MSI2-324 because it is the annotated isoform and, like the more abundant MSI2-346, has the 18-amino-acid insertion generated by the upstream exon 12 acceptor. MSI2-328 localized predominantly to the cytoplasm, whereas MSI2-324 localized primarily to the nucleus (Fig 1G-H). The endogenous pattern therefore reflects two differently localized isoforms rather than one protein present in both compartments.

### Two short elements with opposing effects specify MSI2 localization

MSI2-328 has been the subject of nearly all published work on human MSI2, yet the sequences responsible for its subcellular localization have not been characterized. MSI2 contains two N-terminal RNA recognition motifs (RRMs) and a large C-terminal intrinsically disordered region (IDR) (Fig. 2A). To investigate whether the IDR contributes to cytoplasmic localization, we imaged a panel of GFP-tagged truncations and internal deletions in the MSI2-328 backbone (Fig. 2B). Removing everything downstream of residue 204 (ΔIDR++) did not alter cytoplasmic localization, indicating that this portion of the IDR is not required for MSI2-328 localization. Residues 194–234 are conserved and homologous to the PABPC1-binding region of MSI1 ^23,37^, and deleting them relocalized MSI2-328 to the nucleus (Fig. 2C; Supplementary Fig. S2A). We refer to this region as the cytoplasmic localization region (CLR). Cytoplasmic localization of MSI2-328 therefore depends on a short internal element rather than on the bulk of the disordered region.

**Figure 2.**
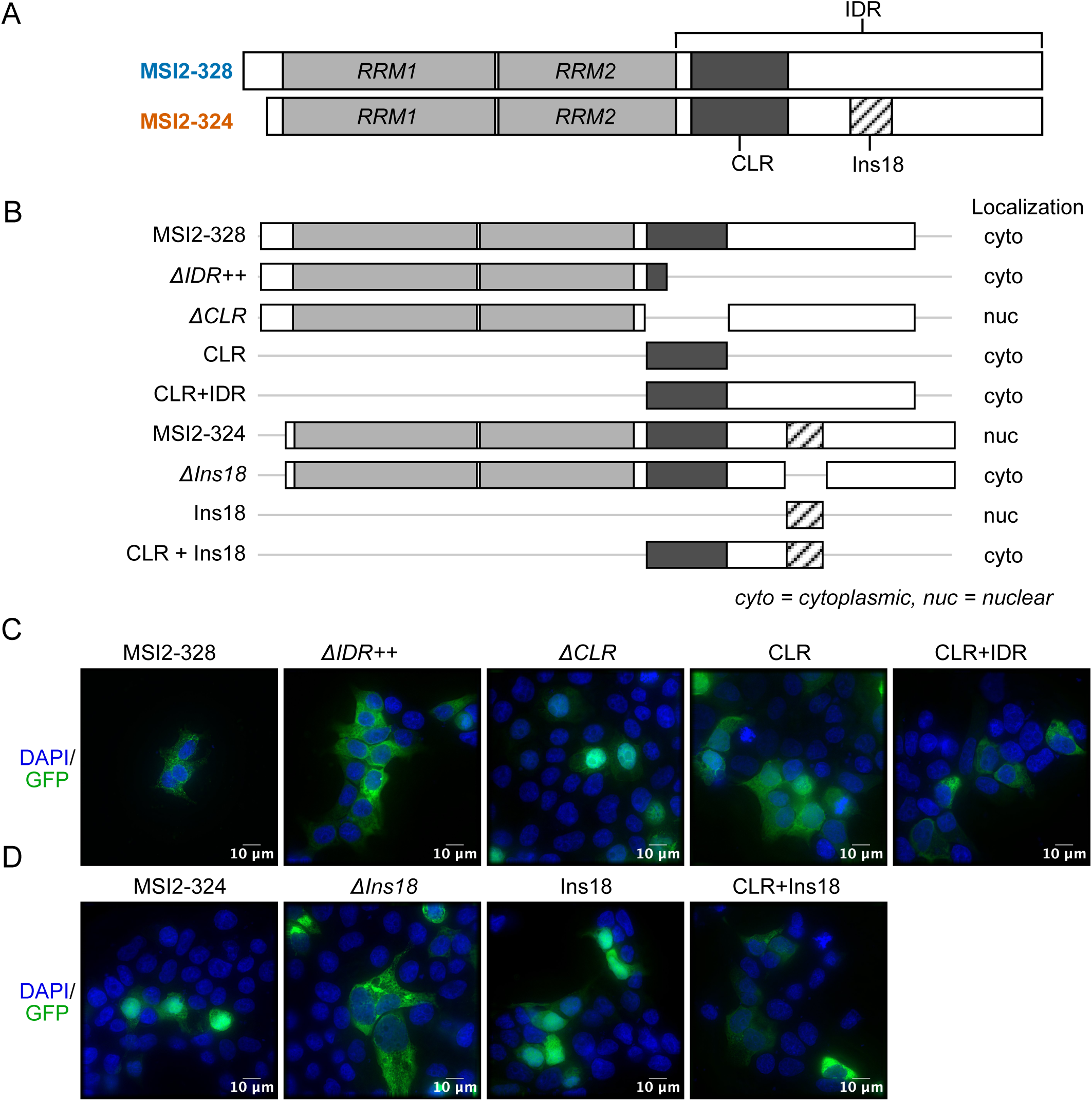
Two short sequence elements with opposing effects specify MSI2 localization. (A) Schematic of MSI2-328 and MSI2-324 showing the RNA recognition motifs (RRM1 and RRM2), cytoplasmic localization region (CLR), 18-amino-acid insertion (Ins18), and intrinsically disordered region (IDR). (B) Schematic of λN-GFP-tagged MSI2 constructs with their observed subcellular localization indicated (C, cytoplasmic; N, nuclear). (C) Representative immunofluorescence of λN-GFP-tagged constructs; MSI2-328, ΔCLR, ΔIDR++, CLR, and CLR+IDR (green) with DAPI (blue). (D) Representative immunofluorescence of λN-GFP-tagged constructs; MSI2-324, ΔIns18, Ins18, and CLR+Ins18 (green) with DAPI (blue). Scale bars, 10 μm.

To test whether the CLR was sufficient to direct cytoplasmic localization, we imaged it alone (CLR) and together with the remaining C-terminal IDR (CLR+IDR). Both localized predominantly to the cytoplasm (Fig 2C, Supp Fig 2A). The CLR is therefore necessary and sufficient for cytoplasmic localization of MSI2-328.

MSI2-324 contains the same CLR as MSI2-328 but localizes predominantly to the nucleus, so we tested whether the 18 aa added by alternative splicing are responsible for this difference. Removing the insertion (ΔIns18) produced cytoplasmic localization comparable to MSI2-328 (Fig 2C, D, Supp Fig 2A-B). The isolated sequence (Ins18) localized predominantly to the nucleus (Fig 2D, Supp Fig 2B). Thus Ins18 is necessary for nuclear localization of MSI2-324 and sufficient on its own.

To examine how these opposing localization elements function together, we tested a construct containing both the CLR and Ins18 (CLR+Ins18). Despite the presence of the insertion, CLR+Ins18 localized predominantly to the cytoplasm (Fig. 2D, Supp Fig 2B). This contrasts with full-length MSI2-324, which contains both regions but remains predominantly nuclear, indicating that the balance between these elements depends on the surrounding protein context.

### Cytoplasmic localization is required for MSI2-mediated translational activation

To measure the regulatory activity of each isoform directly, independent of binding affinity, binding position, and competition with other RNA-binding proteins ^38^, we recruited each isoform to a reporter mRNA through a heterologous high-affinity interaction rather than through its own binding sites. In this tethering assay ^39^, MSI2-328 is fused to the λN peptide, which binds BoxB stem-loops in the 3’ UTR of a nano luciferase (nLuc) reporter co-expressed with firefly luciferase from a bidirectional promoter (Fig 3A). Each construct carries the same N-terminal GFP fusion used for imaging, so localization and translational activity were measured on the same proteins. Tethered MSI2-328 increased reporter activity approximately 7.5-fold relative to tethered GFP (p = 9.3 × 10-4). Untethered GFP and untethered MSI2-328 produced only modest increases of approximately 1.3-fold and 2.1-fold (Fig 3B). Reporter mRNA levels did not differ across conditions (Supplemental Fig 3A), so the increase reflects translation rather than accumulation of the reporter mRNA. Depletion of endogenous MSI2 reduced reporter activity by 49% and 41% for reporters containing synthetic *Numb*-derived or *SCD* 3′ UTR MSI2-binding elements, respectively (Fig. 3C) ^33,40^. These results indicate that MSI2-328 promotes target mRNA translation.

**Figure 3.**
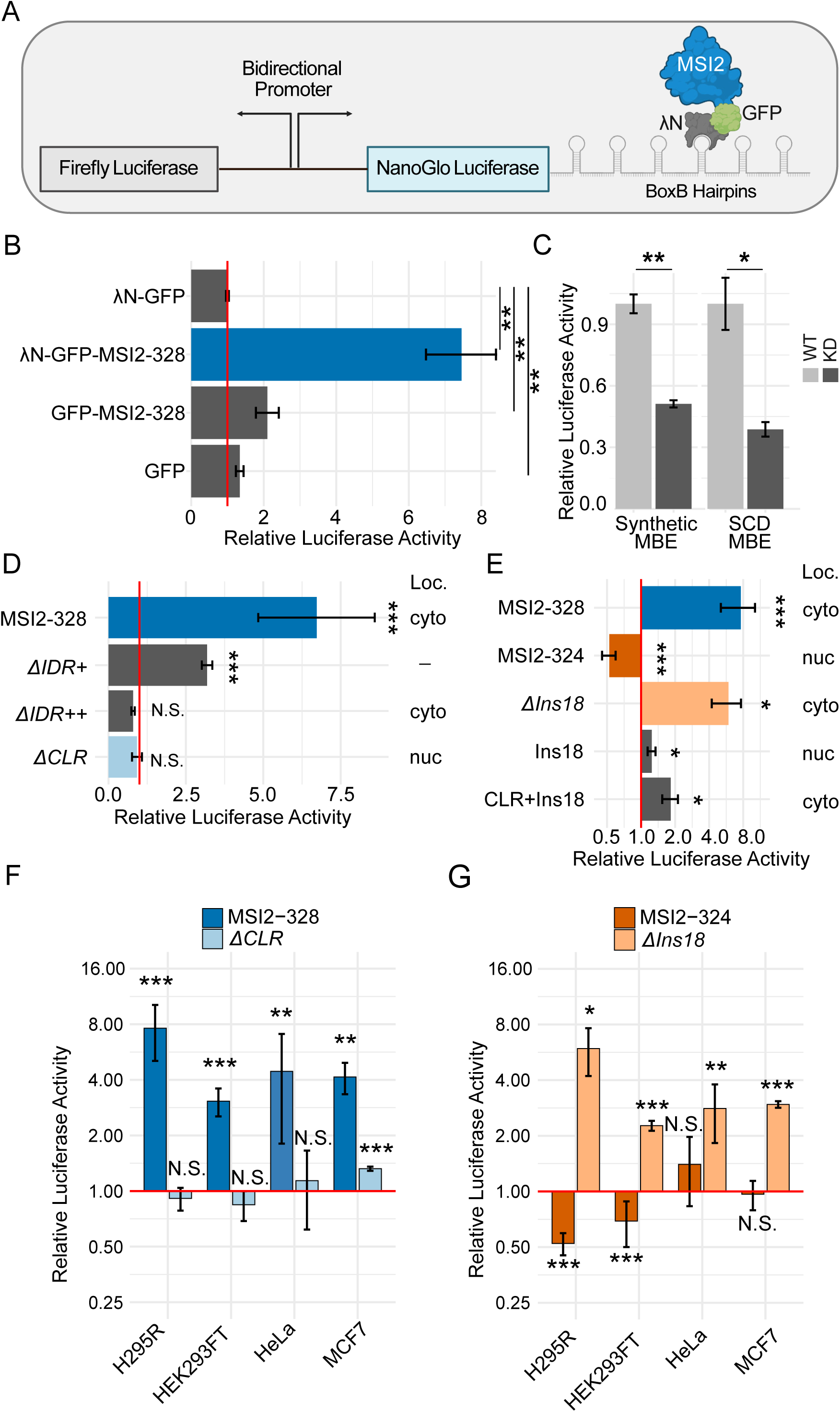
Cytoplasmic localization is required for MSI2-mediated translational activation. (A) Schematic of the tethering-based luciferase reporter assay. (B) NanoLuc/firefly luciferase ratios for the indicated constructs in H295R cells (n = 6), normalized to λN-GFP (red line). (C) Relative luciferase activity from reporters containing synthetic or SCD-derived MSI2-binding elements (MBEs) in MSI2 WT and MSI2 KD H295R cells (n = 6), normalized to MSI2 WT. Relative luciferase activity for (D) MSI2-328 and (E) MSI2-324 truncation, deletion, and sufficiency constructs (n = 6), normalized to λN-GFP (red line). Subcellular localization is indicated (C, cytoplasmic; N, nuclear). (F) Relative luciferase activity from tethering assays with MSI2-328 and MSI2 ΔCLR in the indicated cell lines, normalized to λN-GFP. (G) Relative luciferase activity from tethering assays with MSI2-324 and ΔIns18 in the indicated cell lines, normalized to λN-GFP. Significance was determined using Student’s t-test. (*) P < 0.05; (**) P < 0.01; (***) P < 0.001.

To determine whether the CLR also contributes to translational activation, we tested a series of truncation and deletion constructs using the tethering assay. Disruption of MSI2 RNA binding (ΔRNA) ^36^ or removal of the N-terminal region (ΔN), did not impair translational activation (Supplemental Fig 3B-C). Progressive truncation of the C-terminal IDR reduced MSI2 activity, with ΔIDR+ decreasing activation by 51% and ΔIDR++ completely abolishing it despite robust protein expression (Fig. 3D; Supplementary Fig. S3B–C). Deletion of the CLR (ΔCLR) also abolished activation (Fig 3D; Supplemental Fig S3B-C). Although this region is homologous to the PABPC1-binding domain of MSI1, we found that that MSI2-mediated activation was not strongly dependent on PABPC1 (Supplementary Fig. S3D-F).

Notably, ΔCLR lost both cytoplasmic localization and translational activation, while ΔIDR++ remained cytoplasmic yet was inactive. To determine whether the regions required for activation were sufficient, we examined constructs containing the CLR alone, the distal C-terminal IDR (IDR+), or the extended CLR and IDR (CLR+IDR). Neither the CLR nor IDR+ alone recapitulated activation, and the CLR+IDR produced only a 1.7-fold increase (p = 0.014) (Supplemental Fig S3G-H). The sequence requirements for cytoplasmic localization and translational activation overlap, but cytoplasmic localization alone is not sufficient.

We then compared the regulatory activities of MSI2-328 and MSI2-324 directly. Unlike MSI2-328, tethered MSI2-324 did not activate translation, with reporter activity 48% below tethered GFP (p = 1.2 × 10-10), and reporter mRNA levels unchanged (Fig 3E, Supplemental Fig 3I-J). Removal of the 18aa insertion (ΔIns18) restored cytoplasmic localization and promoted translation 5.9-fold (p = 0.01), comparable to MSI2-328 (Fig 3E). Constructs carrying the insertion alone (Ins18) or together with the shared CLR (CLR+Ins18) produced negligible increases of 1.2-fold (p = 0.017) and 1.8-fold (p = 0.011) (Fig 3E; Supplemental Fig S3K).The insertion therefore abolishes activation in the full-length protein.

Because MSI2-328 and MSI2-324 also differ at their N-termini, we examined whether this difference contributed to their distinct regulatory activities. The ΔIns18 construct retains the alternative N-terminus of MSI2-324 yet showed localization and translational activation comparable to MSI2-328. Similarly, removal of the N-terminal region from MSI2-328 (ΔN) did not impair translational activation (Supplementary Fig. S3B-C). These findings indicate that the 18-amino-acid insertion, rather than the N-terminal difference, accounts for the regulatory differences between MSI2-328 and MSI2-324.

To test whether these results hold across cellular contexts, we repeated the tethering assays in additional cell lines. MSI2-328 promoted translation in all cell lines tested, and activation required the CLR in each. MSI2-324 did not promote translation and in some lines reduced reporter activity, and removing the insertion shifted it toward an MSI2-328-like phenotype in every line (Fig. 3F,G). The distinct translational activities of MSI2 isoforms are broadly conserved across the cellular contexts.

### MSI2 isoforms associate with different cofactors and bind different transcript regions

To determine whether the isoforms bind different protein partners, we performed FLAG immunoprecipitation with RNase treatment followed by mass spectrometry from cells expressing FLAG-MSI2-328 or FLAG-MSI2-324 (Supplemental Fig 4A). Because many proteins, including splicing and hnRNP factors, were recovered with both isoforms, we focused on differences in relative enrichment rather than interactions exclusive to either isoform. MSI2-328 was enriched for translation initiation factors and ribosomal proteins, among them eIF3g, eIF3d, PABPC4, and multiple ribosomal structural proteins (Fig 4A, Supplemental Table 1). MSI2-324 was enriched for nuclear RNA processing factors, among them the U2 snRNP-associated proteins SF3A2 and SF3A3 (Fig 4A, Supplemental Table 1). These differences suggest that the two isoforms associate preferentially with protein networks involved in distinct regulatory processes, consistent with their respective subcellular localization patterns.

**Figure 4.**
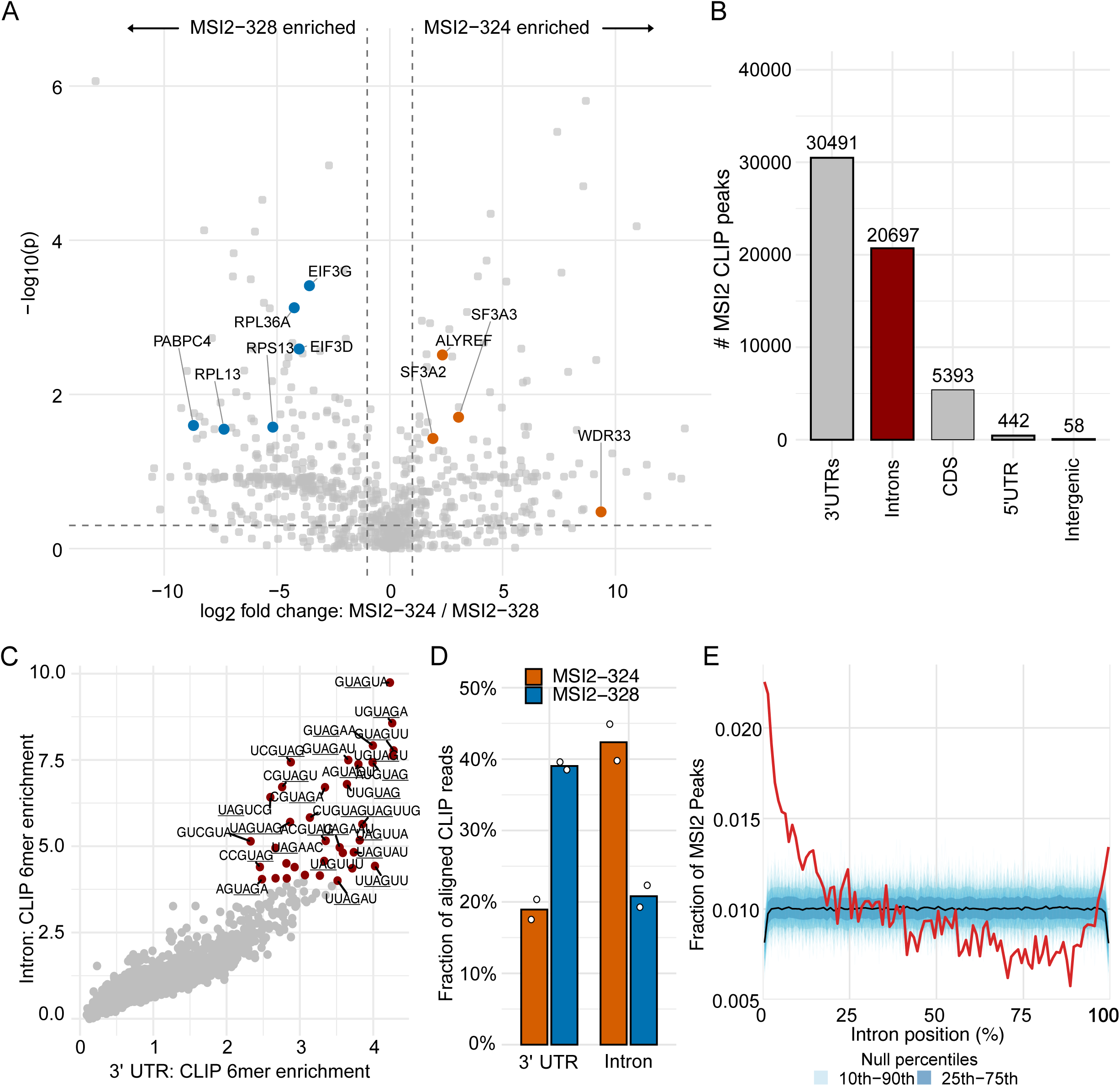
MSI2 isoforms associate with distinct proteins and bind distinct transcript regions. (A) Volcano plot comparing protein enrichment between FLAG-MSI2-324 and FLAG-MSI2-328. Significance thresholds were P < 0.05 and |log2FC| > 1. Translation-associated proteins enriched with MSI2-328 are highlighted in blue; nuclear RNA processing proteins enriched with MSI2-324 are highlighted in orange. (B) Barplot of the number of endogenous MSI2 CLIP-seq peaks across annotated transcript regions. (C) Enrichment of 6mers in endogenous MSI2 CLIP-seq peaks versus background for intronic (y-axis) and 3’ UTR (x-axis) binding sites. 6mers in red were enriched at least 4-fold in either region. (D) Barplot of the fraction of MSI2-324 and MSI2-328 CLIP reads between 3’ UTRs and introns. Bars show mean annotation fractions across two biological replicates per isoform; open circles indicate individual replicates. (E) Distribution of 20,364 endogenous MSI2 CLIP-seq peaks across 9,565 introns. Peak positions were normalized to intron length and grouped into 1% bins, with each intron contributing equal total weight. The red curve shows the observed distribution, cyan curves show 500 size-matched randomized null profiles, and the black curve shows the null median. Light and dark blue bands indicate the 10th-90th and 25th-75th null percentiles.

The association of MSI2 isoforms with cytoplasmic translation factors and nuclear RNA processing proteins raised the possibility that MSI2 also interacts with RNA substrates in both compartments. We performed CLIP-seq on endogenous MSI2 to characterize its RNA-binding sites (Supplemental Fig. 4B-C). As expected from established roles in translation and mRNA stability, 3’ UTRs contained the largest number of binding sites, with 30,491 peaks. Introns contained the second largest, with 20,697 peaks, substantially more than coding sequence, with 5,393 (Fig 4B). These findings suggest that endogenous MSI2 interacts with both mature mRNAs and nuclear pre-mRNA substrates.

To determine whether intronic binding reflects the same sequence preference MSI2 exhibits in 3’ UTRs, we calculated the enrichment of each 6mer within MSI2 peaks relative to its frequency in the corresponding transcript region. Enrichment values in introns and 3’ UTRs were strongly correlated, and the 6mers enriched at least 4-fold in either region all contained UAG (Fig 4C). The difference between intronic and 3’ UTR binding therefore reflects the substrates available to MSI2 rather than a change in its specificity.

To determine whether MSI2 isoforms preferentially bind different transcript regions, we performed CLIP-seq on each isoform individually (Supplemental Fig. 4D-E). MSI2-328 reads mapped predominantly to 3’ UTRs, which accounted for 39% of mapped reads compared to 19% for MSI2-324. Conversely, intronic reads accounted for 42% of MSI2-324 reads compared to 21% for MSI2-328 (Fig 4D; Supplemental Fig. S4F). The size of the shift is consistent with the partial, rather than absolute, compartmentalization we observe by microscopy.

To determine where within introns MSI2 binds, we compared the positions of 20,364 endogenous MSI2 peaks distributed over 9,565 introns against randomized null profiles. Binding was non-uniform, with enrichment toward the 5’ end of the intron (Fig 4E). Indeed, 6.5% of intronic peaks fell within 320 nucleotides of a 5’ splice site, 1.4-fold more than expected from a uniform distribution within the same introns (p < 0.001, 1,000 permutations). Taken together, MSI2 isoforms bind the same motif in different places, and nuclear MSI2-324 is biased to pre-mRNA.

### Usage of the exon 12 acceptor varies across tissues and increases in tumors

The two isoforms can account for context-dependent activity only if their relative abundance varies, so we quantified split reads spanning the exon 11 donor across 19,788 GTEx samples, restricting analysis to the 10,765 samples with at least 20 spanning reads. This approach directly measures usage of the alternative exon 12 splice acceptor responsible for the 18-amino-acid insertion, independently of existing isoform annotations. Usage varied nearly threefold across tissues, from 19.1% in skeletal muscle to 51.1% in brain (Fig 5A). Within brain, usage was highest in the cerebellar hemisphere at 62.5% (Sup Fig 6A). Among muscle-containing tissues, skeletal muscle was the lowest (Sup Fig 6B).

**Figure 5.**
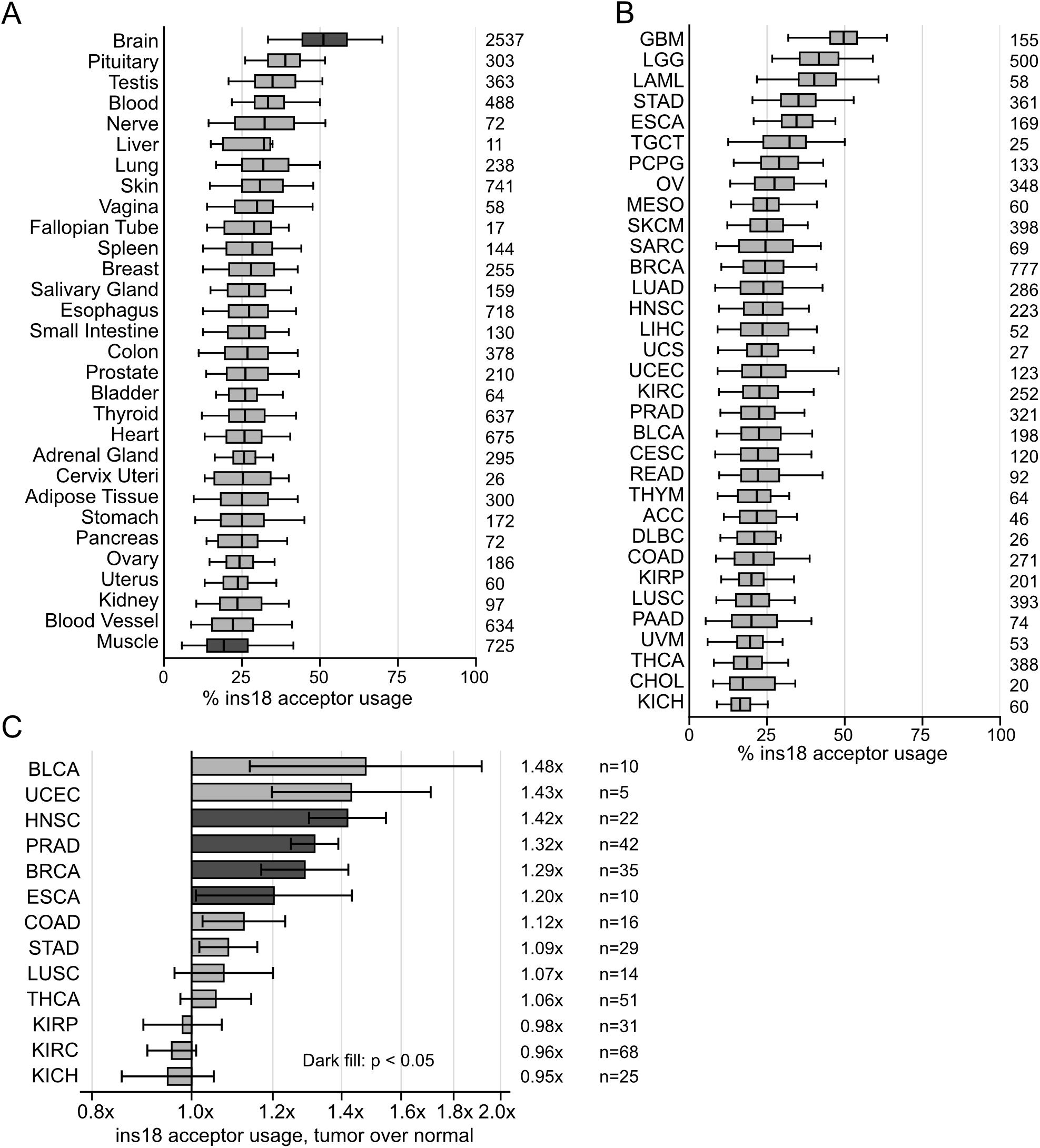
Usage of the exon 12 acceptor varies across human tissues and increases in tumors. (A) Percent of split reads at the two MSI2 exon 12 acceptors that use the insertion-generating acceptor, across GTEx tissue types. Samples with fewer than 20 reads spanning the exon 11 donor were excluded (10,765 of 19,788 samples retained). Bars show medians; error bars show the interquartile range. (B) Acceptor usage across TCGA cancer cohorts. Each point represents an individual tumor sample; center lines indicate medians, boxes represent the interquartile range, and whiskers extend to 1.5X the interquartile range. (C) Fold change in median acceptor usage between tumor and matched normal samples for the 13 TCGA cohorts with at least five matched normals, sorted from largest increase to largest decrease. Error bars are bootstrap 95% confidence intervals. Bars with a bold black outline are significant by Mann-Whitney test (P < 0.05, uncorrected).

We next applied the same analysis to TCGA tumor samples and their matched normals. Tumor samples showed variability across cancer types comparable to that across normal tissues (Fig 5B). In the 13 cohorts with at least five matched normal samples, 10 showed increased usage of the MSI2-324-associated acceptor in tumors, with a median increase of 2.8 percentage points (Wilcoxon signed-rank p = 0.0046; Fig 5C). The same increase was apparent in the 291 patients with paired tumor and normal samples, with a median of 1.57 points (p = 0.0065). Four cohorts reached significance individually: prostate adenocarcinoma (p = 8.0 × 10-5), head and neck squamous carcinoma (p = 0.0010), breast carcinoma (p = 0.027), and esophageal carcinoma (p = 0.048). In contrast, three kidney cancer cohorts showed trends toward decreased acceptor usage. Acceptor usage therefore varies across tissues and shifts toward the insertion in several cancers.

## DISCUSSION

Alternative splicing of MSI2 introduces an 18-amino-acid sequence that alters its subcellular localization and regulatory activity. In this study, we demonstrate that MSI2-328 localizes predominantly to the cytoplasm and promotes translation, whereas MSI2-324 localizes primarily to the nucleus and exhibits distinct RNA-binding and protein interaction profiles. These findings suggest that differences previously attributed to context-specific MSI2 regulation may reflect variation in the relative abundance of functionally distinct isoforms. Many previous functional studies have relied on MSI2 knockdown approaches that deplete multiple isoforms simultaneously ^6,18,41^, while overexpression studies have used the 328-residue isoform alone ^1,18,42,43^. As a result, the contributions of individual MSI2 isoforms to previously reported regulatory phenotypes remain largely unresolved. Wuebben and colleagues proposed that multiple Msi2 isoforms act together to maintain mouse stem cell-like states ^29^. Our results support the importance of considering multiple isoforms while suggesting that their contributions may arise through distinct subcellular regulatory functions.

We identified two sequence elements that influence MSI2 subcellular localization: a 40-aa region (residues 194–234), termed the cytoplasmic localization region (CLR), and the alternatively spliced 18-aa insertion (Ins18), which promotes nuclear localization. The CLR was necessary and sufficient for cytoplasmic localization of MSI2-328, while Ins18 was necessary for nuclear localization of MSI2-324 and sufficient to promote nuclear localization when expressed alone. Interestingly, a construct containing specifically these regions combined (CLR+Ins18) localized predominantly to the cytoplasm, despite both elements also being present in the predominantly nuclear MSI2-324 protein. This suggests that additional regions or structural features influence how these localization elements function together. We also found that cytoplasmic localization alone is insufficient for MSI2-mediated translational activation. Truncation downstream of residue 204 (ΔIDR++) preserved cytoplasmic localization yet abolished translational activation, while the isolated CLR was unable to promote translation despite directing the protein to the cytoplasm. Therefore, translational activation requires both cytoplasmic localization and IDR sequences beyond residue 204. Whether active nuclear transport, cytoplasmic retention, cofactor binding, or a combination localizes MSI2 remains unknown, as does how localization responds to cellular state.

In the cytoplasm, translational activation does not appear to depend on a single cofactor. Given the homology between the MSI2 CLR and the previously characterized PABPC1-binding region of MSI1, we initially considered whether MSI2 promotes translation through a similar mechanism ^23,37^. Deletion of this region abolished detectable PABPC1 binding, but also resulted in nuclear localization, making it difficult to distinguish whether the loss of interaction reflects disruption of a binding interface or reduced access to cytoplasmic PABPC1 (Supplementary Fig. 3D). To further investigate the contribution of PABPC1, we depleted it using two independent siRNAs and observed only modest effects on MSI2-mediated translational activation (Supplementary Fig. 3E-F). We also did not detect GLD2 interactions which were previously proposed to mediate MSI1-dependent translational activation ^24^. Instead, our proteomics analysis identified associations between MSI2-328 and several translation initiation factors and ribosomal proteins. These observations point to broader translation-associated complexes rather than to one dominant cofactor.

Previous studies have implicated Musashi proteins in alternative splicing regulation, particularly through binding to intronic sequences downstream of alternative exons. In mouse photoreceptors, UAG clusters in that region are required for Musashi-dependent inclusion, and at *Ttc8* exon 2A the functional sites lie within ∼320 nucleotides of the 5’ splice site ^26^. Similarly, Musashi CLIP paired with Msi1/Msi2 double-knockout RNA-seq found binding in the same region for 165 exons that lost inclusion upon Musashi loss ^25^. Our results are consistent with these observations, as endogenous human MSI2 preferentially binds UAG-containing sequences and exhibits enrichment near the 5′ ends of introns. Specifically, 6.5% of intronic MSI2 peaks occurred within 320 nucleotides of a 5′ splice site, representing a 1.4-fold enrichment over the expected distribution. Furthermore, isoform-specific CLIP-seq revealed that MSI2-324 preferentially binds intronic regions. However, these published studies establish a splicing function for the Musashi family rather than for MSI2 specifically. The enrichment of MSI2 binding near 5’ splice sites could also reflect differences in the availability of intronic sequences during transcription, rather than recruitment specifically associated with splicing. Alternative splicing of MSI2 therefore influences not only its localization, but also its access to pre-mRNA.

The differential expression of MSI2 isoforms has implications for cancer biology, where MSI2 has been assigned both oncogenic and tumor-suppressive functions. Because the isoforms differ in regulatory function, a shift in their relative abundance could alter MSI2-dependent programs without any change in total MSI2 expression. We observe such a shift in several tumor types, both across cohorts and within individual patients. Earlier work in triple-negative breast cancer likewise found isoform-specific expression differences and reported that one isoform conferred a protective phenotype on overexpression ^44^. Although the factors controlling exon 12 splice acceptor selection remain unknown, identifying them could help explain how MSI2 isoform balance changes across tissues and tumors. Determining whether these changes contribute directly to tumor progression or therapeutic response will require approaches that specifically manipulate the alternative splicing event, rather than broadly inhibiting or depleting MSI2.

Although each step of this model is supported, our conclusions are limited in three ways. First, the tethering reporter measures translational output rather than exon inclusion, so we have not directly tested whether MSI2-324 regulates splicing, though several studies implicate Musashi proteins in splicing ^25,26^. Second, our insertion-containing constructs are based on MSI2-324, the annotated isoform, rather than on MSI2-346, which our splice-junction and PCR analyses indicate is the more abundant insertion-containing species. The two differ only at the N terminus, which did not affect localization or translational activation in the constructs we tested (Fig 2B,C and Sup Fig 3B). Thus we expect MSI2-346 to behave as MSI2-324 does, although we have not tested the N terminus in an insertion-containing background. Confirming this will require isoform-resolved measurement at the protein level.

One additional MSI2 isoform that warrants further investigation is associated with ENST00000322684, which uses an alternative terminal exon and is predicted to encode a 251-residue protein. This transcript may also predominantly use the canonical first exon, given its strong usage across MSI2 transcripts (Fig 1B,C). Unlike the full-length isoforms, the truncated protein lacks the epitope recognized by our C-terminal antibody and would therefore escape detection in both IF and westerns. Usage of this alternative terminal exon appears to be tissue-dependent, accounting for a median of 30% of reads at that splice site in most regions of the brain versus 4.5% in other tissues. The protein retains the CLR but lacks everything beyond residue 242, including the Ins18 region. Our truncation construct ΔIDR+ (residues 1-258), which approximates that architecture, promoted translation at roughly half the level of the full-length MSI2-328. These observations suggest that the truncated isoform may be predominantly cytoplasmic and retain some translational activity, although its localization and function remain to be tested directly.

In summary, one alternative splicing event controls MSI2 subcellular localization, which determines its regulatory function. MSI2-328 localizes predominantly to the cytoplasm, where it binds mature mRNAs and translation-associated proteins and promotes translation. In contrast, the insertion-containing MSI2-324 localizes primarily to the nucleus and preferentially binds intronic sequences, consistent with a role in splicing. Both isoforms recognize the same motif, so what differs between them is which RNAs they encounter rather than which sequences they bind. Alternative splicing also directs QKI and FMRP isoforms to different compartments ^27,45^. While no comparable systematic analysis exists for RBPs, a survey of 693 transcription factor (TF) isoforms from 246 genes found that approximately two-thirds differed from their reference isoform in at least one molecular activity ^46^. Both the ENCODE RBP and TF resources were built at gene resolution, since neither the antibodies used for eCLIP and ChIP-seq nor the shRNAs are designed to distinguish isoforms ^47–49^. Altogether, a single reference isoform does not capture the functional diversity of a gene. Isoform usage varies across tissues and shifts in disease, so resolving individual isoforms, in studies of single proteins as much as in large-scale surveys, is what will determine how often alternative splicing changes where proteins act rather than what they bind.

## MATERIALS AND METHODS

### Construct cloning

MSI2-328 and MSI2-324 constructs were derived from the annotated transcripts ENST00000284073 and ENST00000416426, respectively. FLAG-MSI2 sequences were synthesized by Twist Bioscience and cloned into the pRD-RIPE vector for doxycycline-inducible expression as described ^33^. A bidirectional luciferase reporter plasmid containing NanoLuc and firefly luciferase was synthesized by Twist Bioscience. The plasmid was linearized with HindIII, and sequences containing 15 BoxB hairpins ^50^, a 390-nt SCD MBE encompassing the highest-confidence MSI2 CLIP-seq peak, or seven synthetic MSI2 binding sites (Syn;^33^) were inserted into the NanoLuc 3′ UTR by Gibson assembly. Tethering constructs were generated by Gibson assembly of MSI2 sequences synthesized by Twist Bioscience into the d3_n-EGFP-LacZ backbone ^50^ using primers containing homologous overhangs. Untethered controls were generated by removing the λN sequence from the same backbone. All generated plasmids were verified by whole-plasmid sequencing (Plasmidsaurus). Construct designs, primer sequences, and corresponding protein sequences are provided in Supplementary Table S2.

### Cell culture

H295R cells were cultured in complete media (DMEM/F12, 10% cosmic calf serum, 1% insulin-transferrin-selenium). K562 cells were cultured in RPMI 1640 with L-glutamate and 10% fetal bovine serum. HEK293 and HeLa cells were cultured in DMEM high glucose with 10% fetal bovine serum. MCF7 cells were cultured in DMEM high glucose supplemented with 10% fetal bovine serum, 1% penicillin-streptomycin, and 2 mM L-glutamine. All cells were cultured at 37°C and 5% CO2.

Stable MSI2 knockdown cells were generated by lentiviral transduction using pLV[shRNA]-EGFP:T2A>hMSI2[shRNA#6] (VectorBuilder), targeting the human MSI2 3′ UTR (TRCN0000423713). Cells were transduced at a multiplicity of infection (MOI) of 10 in the presence of 5 µg/mL polybrene and maintained under blasticidin selection.

### Luciferase reporter assay

Cells were plated at 25,000 cells per well in 96-well plates and reverse-transfected with 25 ng of dual-luciferase reporter plasmid containing BoxB or MSI2 binding elements (MBEs) and 25 ng of MSI2 expression plasmid using Lipofectamine 2000 (Thermo Fisher Scientific) according to the manufacturer’s instructions. For RNA and protein collection, cells were plated at 250,000 cells per well in 24-well plates and reverse-transfected with 200 ng each of reporter and expression plasmids. Medium was replaced after 24 h, and cells were harvested 48 h after transfection. Luciferase activity was measured using the Nano-Glo Dual-Luciferase Reporter Assay System (Promega) on a GloMax Navigator (Promega) with dual injection.

For RNA and protein analysis, cells were washed with PBS and lysed in RIPA buffer containing protease inhibitors. Lysates were incubated on ice for 5 min, triturated, and divided equally, with one half transferred to TRIzol LS for RNA extraction and the remainder retained for western blot analysis.

### Microscopy

Cells were seeded into 12-well plates containing 18 mm poly-D-lysine coated German glass coverslips (Fisher Scientific 501929579) and allowed to adhere overnight. Cells were transfected as indicated, washed once with 1X PBS, and fixed in neutral buffered formalin for 10 min at RT followed by washing three times with PBS then blocked and permeabilized in CAS-Block containing 0.2% Triton X-100 (Invitrogen 008120) for 30 min at RT. Primary antibody against MSI2 or FLAG (Abcam ab76148, Sigma F1804) in CAS-T and incubated at 4°C overnight. Samples were washed with PBS containing 0.1% Tween-20 (PBS-T). Secondary antibody (anti-rabbit IgG Fab2 AlexaFluor 647, Cell Signaling 4414) was diluted 1:1000 in PBS-T and incubated for 1 h at RT in the dark, followed by two PBS-T washes. Coverslips were mounted onto glass slides (Corning 2948-75X25) using Vectashield mounting medium containing DAPI (Vector Laboratories H-1800-2). Images were captured on a DeltaVision Ultra with a 60X objective in 1.5000 immersion oil over 20 focal planes of 0.4 μm. All images were acquired with identical laser intensity, focal plane number, and step size. Images were deconvolved and quick projected using SoftWorx (GE Healthcare Life Sciences, version 7.2.2) with maximum intensity projection. Mean intensity ratio quantifications were performed using Fiji/ImageJ (version 2.14.0/1.54p). The nucleus and cytoplasm were outlined manually for each cell. The mean gray value (integrated intensity divided by ROI area) of the fluorescence signal was measured in each compartment and the nuclear/cytoplasmic ratios were calculated for each cell.

### RT-qPCR

RNA was extracted using the Direct-zol RNA MicroPrep or MiniPrep kit (Zymo Research) according to the manufacturer’s instructions. For RNA collected from dual-luciferase reporter experiments, plasmid DNA was depleted as described ^51^. Total RNA (100 ng) was reverse-transcribed using the iScript cDNA Synthesis Kit (Bio-Rad), and RT-qPCR was performed as described ^33^using reporter-specific primers (Supplementary Table S2).

For MSI2 isoform composition experiments, RNA was reverse-transcribed using oligo(dT) primers and Induro Reverse Transcriptase (NEB) according to the manufacturer’s instructions. RT-qPCR primers were designed to distinguish the alternative exon 12 splice acceptors and first exons. Insertion-specific primers amplified transcripts containing the 54-nt exon 12 insertion, while a second primer pair amplified both insertion-containing and insertion-lacking transcripts. Alternative first exon usage was quantified using exon-specific forward primers and a shared reverse primer in exon 2. RPS20 was used as a reference gene. RT-qPCR was performed as described above.

For RT-PCR, forward primers targeting the canonical or alternative first exon were paired with a shared reverse primer within the exon 12 insertion. Products were resolved by agarose gel electrophoresis. Primer sequences are provided in Supplementary Table S2.

Isoform composition was calculated from Cq values using 2^−ΔCq ratios. The insertion-containing fraction was calculated as the insertion-specific signal divided by the total exon 12 signal, and alternative first exon usage as the alternative first exon signal divided by the sum of the canonical and alternative first exon signals.

### Western blot

Cells were collected and western blots performed as described ^33^. Primary antibodies used were MSI2 (Abcam ab76148), GFP (enQuireBio QAB10298), H3 (Abcam ab18521), FLAG M2 (Sigma F1804), and PABP (Abcam ab21060). Signals were detected by chemiluminescence with SuperSignal West Dura Extended Duration Substrate (Thermo Scientific) on an Azure Biosystems Sapphire biomolecular imager.

### PABPC1 siRNA knockdown

HEK293 cells were reverse-transfected with two independent siRNAs targeting PABPC1 (10 pmol per well) or a non-targeting control siRNA, together with 25 ng each of the BoxB luciferase reporter and λN-GFP-MSI2-328 or λN-GFP control constructs in 96-well plates. Transfections were performed using Lipofectamine 2000 as described above. Luciferase activity was measured using the Nano-Glo Dual-Luciferase Reporter Assay System (Promega). PABPC1 knockdown efficiency was assessed by RT-qPCR, with GAPDH used for normalization. siRNA sequences are provided in Supplementary Table S2.

### Immunoprecipitation

H295R cells containing doxycycline-inducible 3×FLAG-MSI2 constructs integrated at a loxP locus were grown on 10- or 15-cm plates to approximately 60% confluence and induced with 2.5 µg/mL doxycycline for 24 h. Immunoprecipitation was performed as described ^33^ with modifications. FLAG-tagged proteins were immunoprecipitated using prewashed anti-FLAG M2 magnetic beads (Sigma M8823) with end-over-end rotation for 3 h at 4°C. Beads were washed four times with NT2 buffer containing RNase I (1.5 U per mg protein input) and resuspended in NT2 buffer. For IP-western analysis, samples were prepared with 4× Laemmli sample buffer containing β-mercaptoethanol. Western blotting was performed as described above using anti-FLAG M2 (Sigma F1804), anti-PABPC1 (Abcam ab21060), and anti-H3 (Abcam ab18521).

For IP-mass spectrometry, immunoprecipitations were performed as described above. Bound proteins were eluted with five packed bead volumes of 0.1 M glycine (pH 3.5) and neutralized with 0.5 M Tris, 1.5 M NaCl (pH 8.0). An aliquot (2.5%) was reserved for IP-western analysis, and the remainder was snap-frozen in liquid nitrogen and submitted to the Fred Hutchinson Cancer Center Proteomics and Metabolomics Core Facility (RRID:SCR_022618) for mass spectrometry.

### CLIP-seq

H295R cells were grown on 15 cm plates in complete media to 60% confluence, washed in 1X PBS, and treated with 300 mJ UV-254 nm (Vari-X-Link) before scraping and pelleting. Three 15 cm plates of cells were used per CLIP-seq replicate.

CLIP-seq was performed as described ^52^ with modifications. Cell pellets were lysed in iCLIP lysis buffer, quantified by BCA, then treated with RNase I at 1.5 and 1.0 U per mg protein lysate input. A 5% size-matched input was taken from the lysate, processed with SP3 beads, and treated in parallel with the CLIP samples following IP. For endogenous MSI2 CLIP-seq, Dynabeads Protein A (Thermo Fisher Scientific) conjugated to anti-MSI2 antibody (Abcam ab76148) were used for immunoprecipitation. For isoform-specific CLIP-seq, H295R cells expressing FLAG-MSI2-328 or FLAG-MSI2-324 were induced with 2.5 µg/mL doxycycline for 24 h before crosslinking and immunoprecipitated using anti-FLAG M2 antibody. All subsequent steps were performed identically. After IP, beads were washed and treated with T4 polynucleotide kinase. Pre-adenylated adapter (/5Phos/AGATCGGAAGAGCACACGTCTG/3Cy55Sp/) was ligated to the 3’ ends of RNP complexes with T4 RNA ligase I (NEB M0437M), followed by free adapter removal with 5’ deadenylase (NEB M0331S) and RecJ endonuclease (Biosearch Technologies RJ411250). After eluting the 3’-ligated RNP complexes into 1X Laemmli buffer, samples were run on a 4-12% Bis-Tris NuPAGE mini protein-gel in 1X MOPS SDS running buffer for 50 min at 180 V. The gel was transferred to a nitrocellulose membrane using the iBlot2 P0 program, then imaged on the Azure Sapphire biomolecular imager for Cy5 signal. Fluorescent RNP complexes from 55 to 100 kDa were excised, treated with Proteinase K, and RNA extracted. Isolated RNA was used as input into the QIAGEN miRNA-seq library prep kit following the manufacturer’s instructions, substituting Superscript IV reverse transcriptase (Thermo Fisher) during cDNA synthesis. Libraries were sequenced on a NovaSeq 6000 for 30 million paired-end reads (2 × 150 bp) each at the University of Colorado Anschutz Medical Campus Cancer Center Genomics Shared Resource Core Facility (RRID:SCR_021984).

### CLIP-seq analysis

Read 1 FASTQ files were used for CLIP-seq analysis. Unique molecular identifiers (UMIs; 12 nt) were extracted using UMI-tools ^53^, identifying a constant adapter sequence (AACTGTAGGCACCATCAAT) immediately upstream of the UMI, allowing up to two mismatches. Low-quality reads were removed using FASTX-Toolkit v0.0.14, requiring at least 95% of bases to have a Phred quality score ≥10. Adapters were trimmed using cutadapt v3.5 ^54^, with a minimum read length of 20 nt and quality trimming thresholds of 15 and 10 at the 5′ and 3′ ends, respectively. Reads were deduplicated using UMIs with custom Python and Perl scripts, and repetitive sequences were removed before alignment to the human reference genome (GRCh38/hg38) using STAR v2.7.9a ^55^, requiring uniquely mapped reads and end-to-end alignment with a maximum mismatch rate of 4%. Endogenous MSI2 CLIP-seq peaks were called using omniCLIP v0.20 ^56^. Genomic annotations of endogenous peaks and isoform-specific aligned reads were determined using custom R scripts. Isoform-specific CLIP-seq data were processed using the same alignment pipeline without peak calling.

Aligned reads from MSI2-328 and MSI2-324 CLIP-seq were assigned to transcript regions based on overlapping genomic annotations, with reads overlapping multiple categories weighted equally among them. Category fractions were calculated relative to all eligible aligned reads for each biological replicate and averaged across two replicates per isoform without size-matched input normalization.

Endogenous MSI2 CLIP-seq peak summits were assigned to overlapping introns longer than 500 nt, yielding 20,364 peaks across 9,565 introns. Summit positions were normalized to intron length and grouped into 100 bins, with each intron contributing equal total weight. Null distributions were generated from 500 randomizations of summit positions within their assigned introns, preserving intron lengths and peak counts while excluding 12.5 nt at each intron end. Observed distributions were compared with the null median and percentile intervals.

### Splice junction analysis

Split reads spanning the shared MSI2 exon 11 donor (chr17:57652162, hg38) were counted for the two annotated 3’ acceptors, at chr17:57674917 (generating the 18 amino acid insertion) and chr17:57674971 (not generating it). Acceptor usage was expressed as the fraction of reads at the two acceptors that use the insertion-generating acceptor. GTEx v11 and TCGA values were computed from STAR junction counts, restricting to samples with at least 20 reads spanning the donor (10,765 of 19,788 GTEx samples). Developmental and cell-type analyses used the Snaptron srav3h compilation ^57^ of public human RNA-seq, restricting to samples with at least 50 spanning reads; samples were classified from SRA run and sample metadata, and per-study values were pooled across samples by summing reads rather than averaging per-sample fractions. Category-level values are medians across samples, with 95% confidence intervals obtained by bootstrap resampling of samples (10,000 resamples). Paired comparisons across studies used the Wilcoxon signed-rank test.

### Tumor and normal comparisons

TCGA tumor samples and matched normal samples were analyzed with the same junction-counting procedure. Cohorts were included if at least five matched normal samples passed the read threshold, which retained 13 cohorts. Cohort-level comparisons used the Mann-Whitney test without correction for multiple comparisons, because the cohorts represent independent tumor types rather than repeated tests of one hypothesis. The paired analysis was restricted to the 291 patients from whom both a tumor and a normal sample were available and used the Wilcoxon signed-rank test.

## Data and Code Availability

Raw and processed CLIP-seq data generated in this study have been deposited in the Gene Expression Omnibus (GEO) under accession GSE352611. Custom scripts used for CLIP-seq processing, genomic annotation, splice junction analysis, and figure generation are available at GitHub https://github.com/mukherjeelab/2026_MSI2_isoforms.

## ACKNOWLEDGMENTS

This material is based upon work supported by the National Science Foundation Graduate Research Fellowship Program under Grant No. 1000317291-NSFGRFP-Walters (K.W.), the Research Experiences for Graduate and Medical Students (REGMS) Summer Fellowship Program (K.W.), the Victor W. Bolie and Earleen D. Bolie Graduate Scholarship Fund (K.W.), the American Cancer Society Postdoctoral Fellowship (PF-24-1325387-01-RMC) (N.K.), Polish National Agency for Academic Exchange Bekker Program PPN/BEK/2019/1/00173 (M.P.S.), the University of Colorado Anschutz Medical Campus RNA Bioscience Initiative (N.M., M.P.S), the Boettcher Foundation Webb-Waring Early Career Investigator Award AWD-103075 (N.M.), and the National Institutes of Health 1R35GM147025-01 (N.M.). BioRender was used in Figure 3A.

## AUTHOR CONTRIBUTIONS

N.M. conceived the project; K.W., N.K., A.B. and R.T. performed experiments and collected data; K.W., N.K., M.P.S., and N.M. performed formal analysis and conducted the visualization; K.W. and N.M. wrote the original draft; K.W., N.K., A.B., R.T., M.P.S., and N.M. reviewed and edited the paper; K.W., N.K., M.P.S., and N.M. acquired funding; N.M. provided resources; N.M. supervised the project.

## SUPPLEMENTAL FIGURE LEGENDS

**Supplemental Figure S1. MSI2 transcript isoforms and alternative splice site usage across GTEx samples.** (A) Schematic representation of annotated, protein-coding MSI2 splice isoforms (MSI2-328, MSI2-324, MSI2-251) with their corresponding Ensembl transcript IDs and amino acid (aa) length. Exons are represented as black boxes, while isoform specific regions are highlighted in orange or gray. (B) Distribution of alternative first exon usage (top) and exon 12 alternative splice acceptor usage (bottom) across GTEx samples. Histograms show the percentage of junction-spanning reads supporting each alternative splice event per sample. Sample sizes, median usage percentages, and aggregate junction read counts are indicated.

**Supplemental Figure S2. Subcellular localization of MSI2 deletion, truncation and sufficiency constructs.** (A) Representative immunofluorescence single-channel and merged images of GFP-tagged constructs; MSI2-328, ΔIDR++, ΔCLR, CLR, and CLR+IDR (green) with DAPI (blue). (B) Representative immunofluorescence of single channel and merged images of GFP-tagged constructs; MSI2-324, ΔIns18, Ins18, and CLR+Ins18 (green) with DAPI (blue). Scale bars, 10 μm.

**Supplemental Figure S3. Characterization of MSI2 translational activity and functional domains.** (A) Relative NanoLuc reporter mRNA levels measured by RT-qPCR (n = 3), normalized to λN-GFP (red line). (B) NanoLuc/firefly ratios from tethering assays with MSI2-328, truncation constructs (ΔN, ΔIDR+, ΔIDR++, ΔCLR), RNA binding mutant (ΔRNA), and untethered controls (NoTether). (C) Western blot probing GFP to assess expression of MSI2 mutant tethering constructs in HEK293 cells. H3 is used as a loading control. (D) FLAG immunoprecipitation followed by western blot for endogenous PABPC1. Input (0.5%) is shown on the left and immunoprecipitated samples (50%) on the right. (E) Relative luciferase activity in HEK293 cells following PABPC1 knockdown with two independent siRNAs (n = 4), normalized to λN-GFP (red line). (F) Relative PABPC1 transcript expression normalized to scramble siRNA (n = 1). (G) Relative luciferase activity for MSI2-328 sufficiency constructs in H295R cells (n = 6), normalized to λN-GFP (red line). (H) Western blot probing GFP to assess expression of MSI2-328 mutant and sufficiency tethering constructs in H295R cells. H3 is used as a loading control. (I) Relative luciferase activity from tethering assays with MSI2-328 and MSI2-324 in H295R cells, normalized to λN-GFP (red line). (J) Relative NanoLuc reporter mRNA levels for MSI2-328 and MSI2-324 measured by RT-qPCR, normalized to λN-GFP (red line). (K) Western blot probing GFP to assess expression of MSI2-324 sufficiency tethering constructs in H295R cells. H3 is used as a loading control. Significance was determined using Student’s t-test. (*) P < 0.05; (**) P < 0.01; (***) P < 0.001.

**Supplemental Figure S4. Validation and characterization of MSI2 immunoprecipitation and CLIP-seq experiments.** (A) Immunoprecipitation (IP)-Western blot of FLAG-MSI2 constructs from replicate B samples submitted for mass spectrometry. (B) IP-western showing input, unbound supernatant, and 2.5% IP for endogenous MSI2 CLIP-seq samples. (C) Nitrocellulose membrane with MSI2 CLIP-seq and size-matched input samples after ligation to the Cy5.5-containing 3’ adapter. Boxes indicate where RNP complexes were excised. (D) IP-western showing input, and 2.5% IP for FLAG-MSI2-328 and FLAG-MSI2-324 CLIP-seq experiments. (E) Nitrocellulose membrane with isoform specific FLAG-MSI2-328 and FLAG-MSI2-324 CLIP-seq and size-matched input samples after ligation to the Cy5.5-containing 3’ adapter. (F) Distribution of MSI2-324 and MSI2-328 CLIP reads across 3’ UTR, 5’ UTR, CDS, intron, and combined other annotations. Bars show mean annotation fractions across two biological replicates per isoform; open circles indicate individual replicates. Orange, MSI2-328; blue, MSI2-324.

**Supplemental Figure S5. MSI2 exon 12 splice acceptor usage across brain and muscle tissues.** (A) Distribution of exon 12 insertion-containing splice acceptor usage across human brain regions in GTEx samples. (B) Distribution of exon 12 insertion-containing splice acceptor usage across muscle-containing tissues in GTEx samples. Boxplots show the percentage of junction-spanning reads supporting insertion-containing splice acceptor usage. Sample sizes are indicated for each tissue.

