## Supplemental Figure 1 for "Alternative splicing switches human Musashi-2 between cytoplasmic and nuclear regulatory functions"

Sup Figure 1

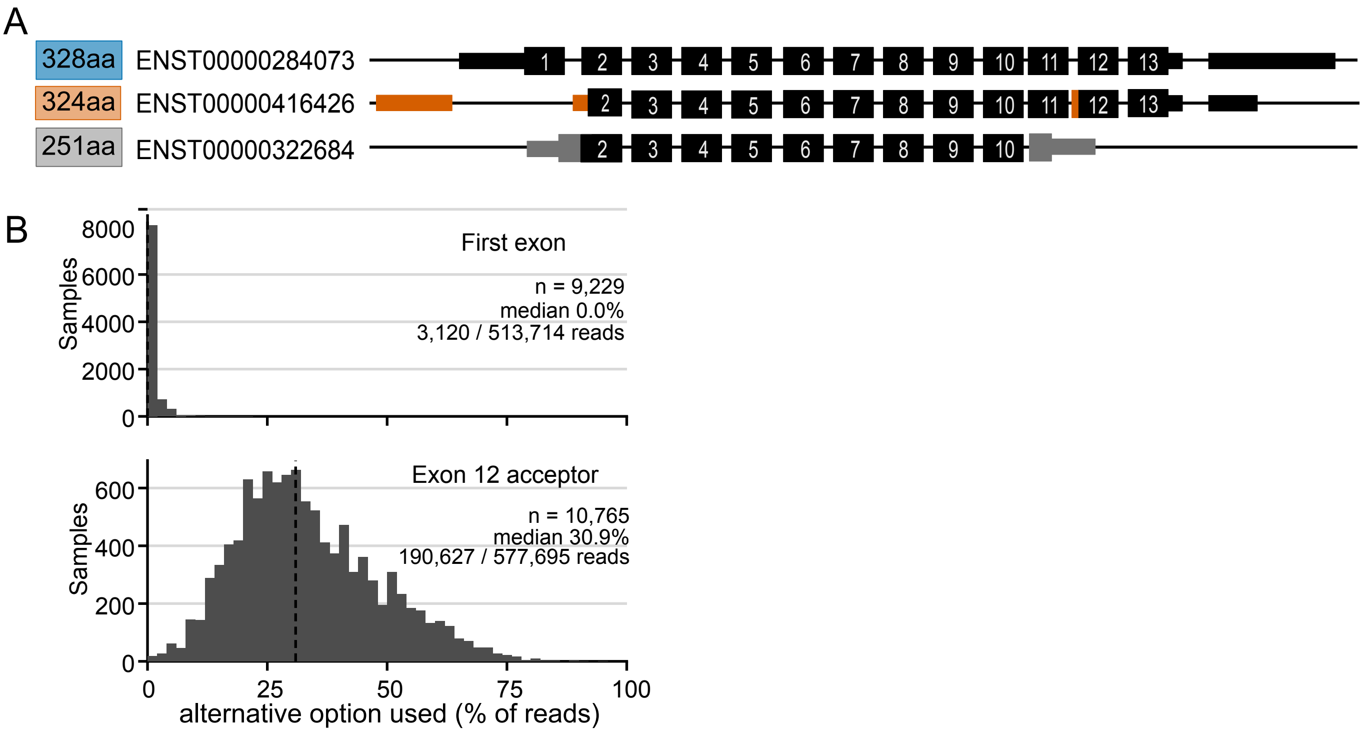

A
328aa
ENST00000284073
324aa
ENST00000416426
251aa
ENST00000322684
1
2
3
4
5
6
7
8
9
10
11
12
13
B
First exon
n = 9,229
median 0.0%
3,120 / 513,714 reads
Samples
8000
6000
4000
2000
0
Exon 12 acceptor
n = 10,765
median 30.9%
190,627 / 577,695 reads
600
400
200
0
25
50
75
100
alternative option used (% of reads)
