## Supplementary figures and images for "Alternative splicing switches human Musashi-2 between cytoplasmic and nuclear regulatory functions"

### Supplemental Figure 2

Supp Figure 2

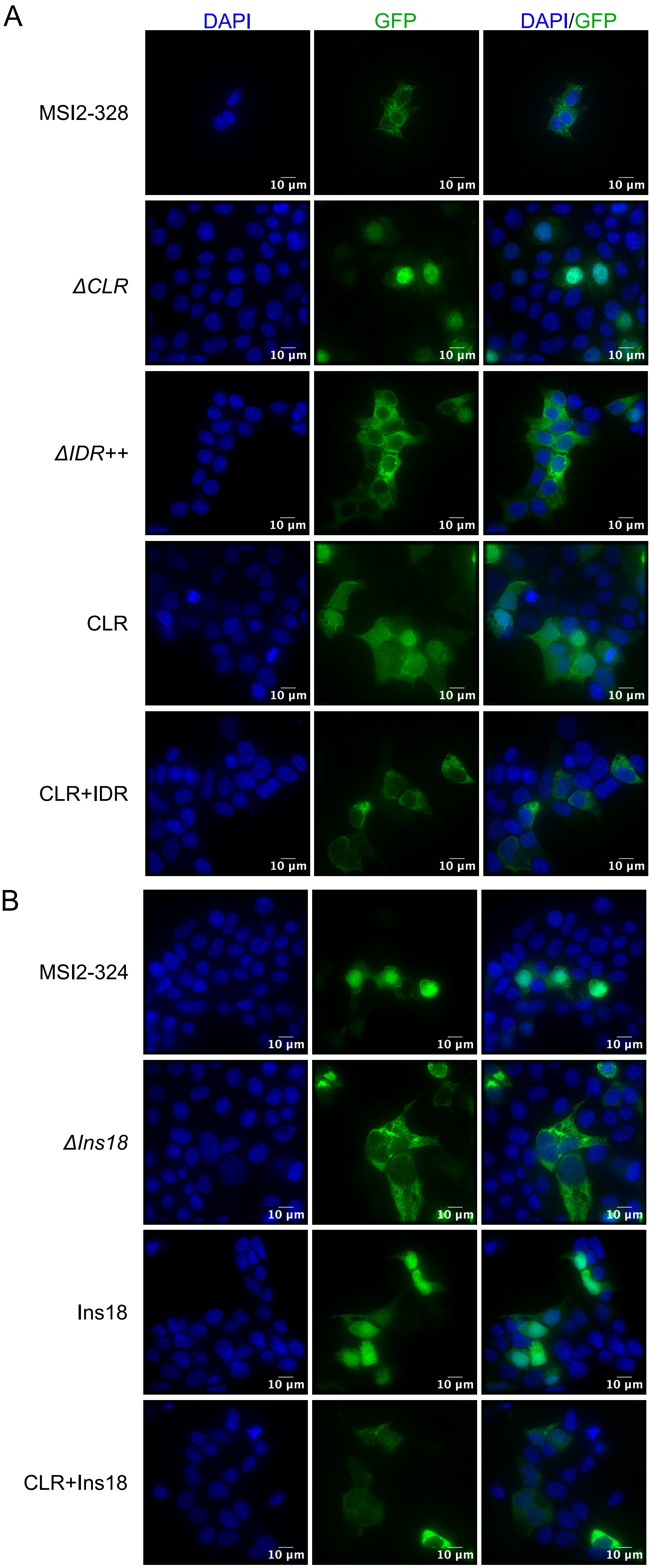

### Supplemental Figure 3

Supp Figure 3

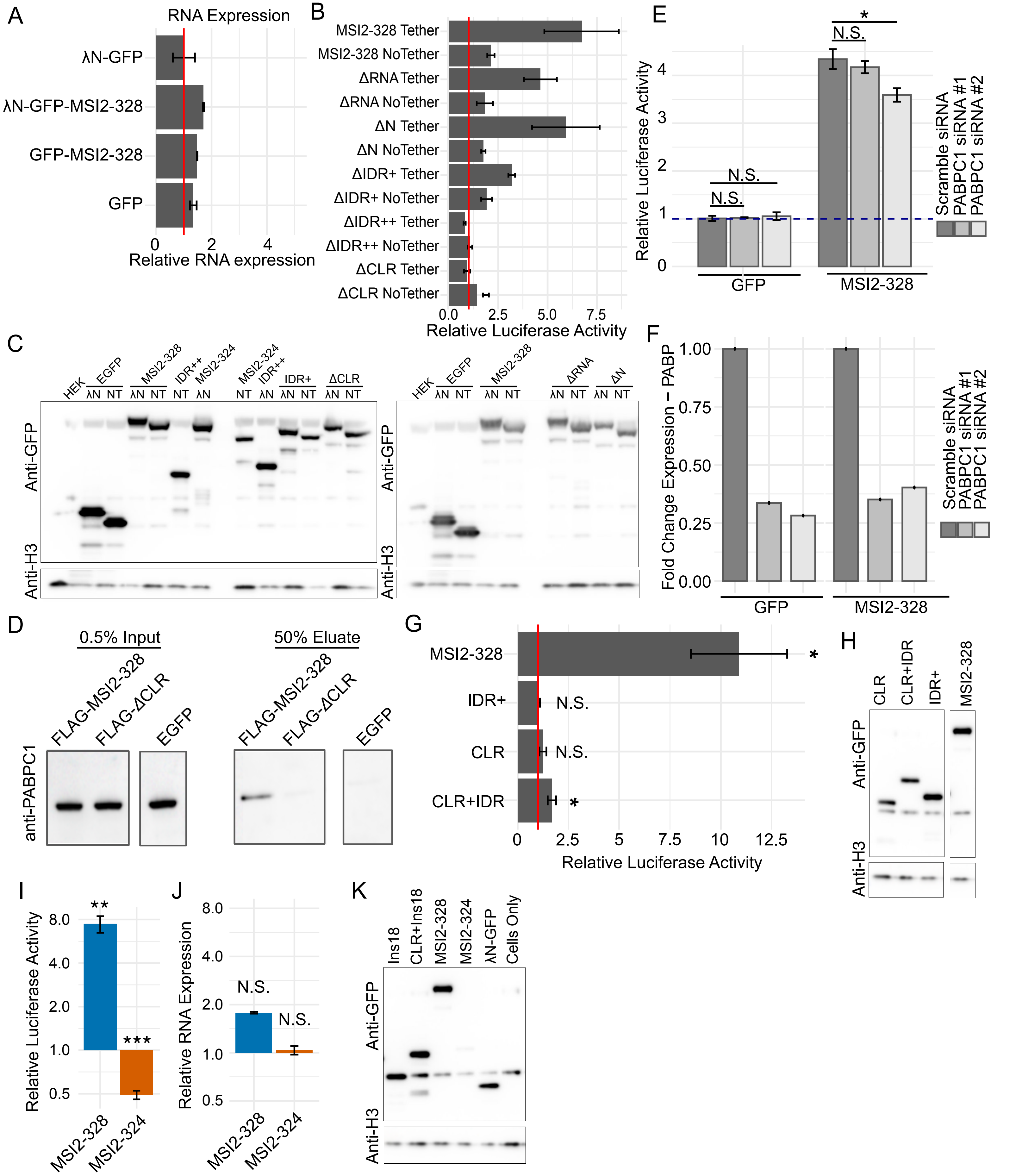

### Supplemental Figure 4

Supp Figure 4

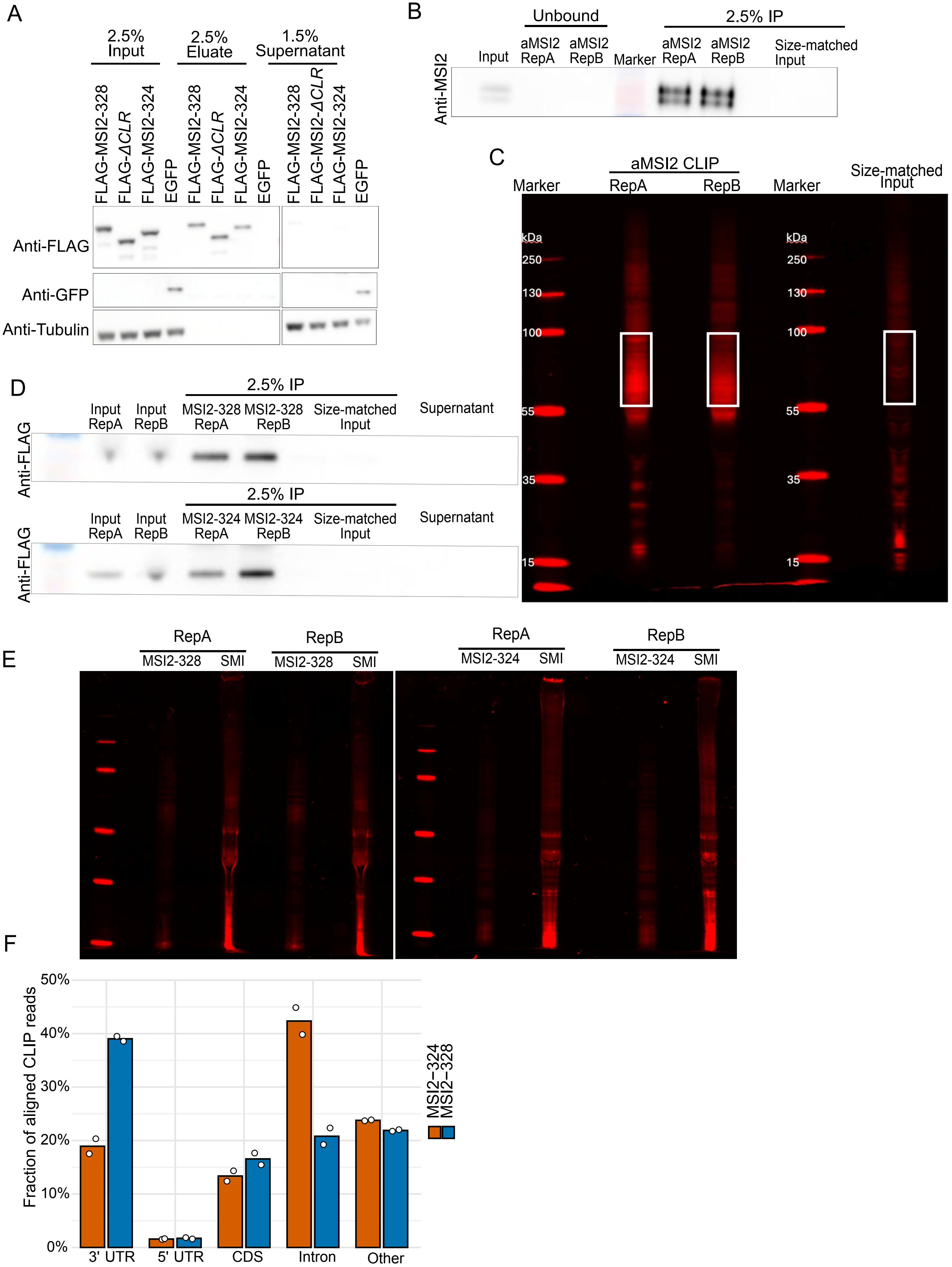
