## Supplemental Figure 5 for "Alternative splicing switches human Musashi-2 between cytoplasmic and nuclear regulatory functions"

Supp Fig 5

A

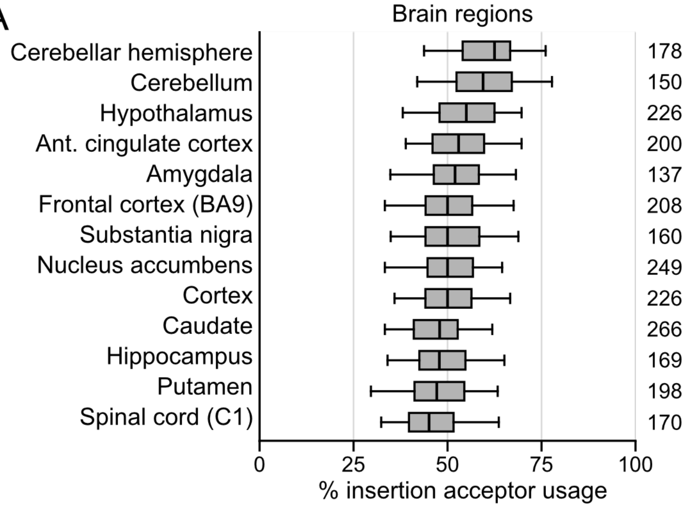
Brain regions
Cerebellar hemisphere
Cerebellum
Hypothalamus
Ant. cingulate cortex
Amygdala
Frontal cortex (BA9)
Substantia nigra
Nucleus accumbens
Cortex
Caudate
Hippocampus
Putamen
Spinal cord (C1)
178
150
226
200
137
208
160
249
226
266
169
198
170
0
25
50
75
100
% insertion acceptor usage

B

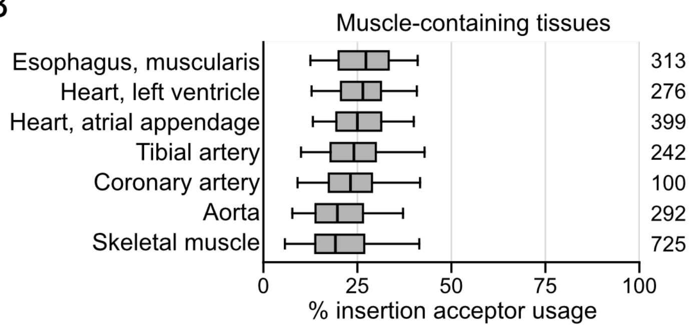
Muscle-containing tissues
Esophagus, muscularis
Heart, left ventricle
Heart, atrial appendage
Tibial artery
Coronary artery
Aorta
Skeletal muscle
313
276
399
242
100
292
725
0
25
50
75
100
% insertion acceptor usage
