## Supplemental Methods for "Alternative splicing switches human Musashi-2 between cytoplasmic and nuclear regulatory functions"

MSI2 Paper Supplemental Methods

**Construct cloning**

MSI2 constructs were ordered from Twist Biosciences Corporation with a 5' flank sequence of CAGGTGTCGGTACCGCGGGCCCGGGATCCA and a 3' flank sequence of CTGGCTCACAAATACCACTGAGATC for each construct. The pRD-RIPE plasmid was digested with AgeI and BstXI. Each MSI2 gene fragment was amplified by PCR with NEB Q5 high fidelity 2X mastermix and primers containing homology arms to the pRD-RIPE vector. The amplified inserts and digested vector were combined with NEBuilder HiFi DNA assembly master mix for Gibson Assembly. Resulting assemblies were transformed into NEB Stable Competent cells and screened for positive colonies by PCR. Positive clones were grown overnight at 30°C and plasmid isolated by ZYMO Research plasmid miniprep classic kit. Whole plasmid sequencing was performed by Plasmidsaurus. The dual luciferase plasmid was cloned as described (Walters et al. 2023).

**Cell culture**

H295R cells were cultured in complete media (DMEM/F12, 10% cosmic calf serum, 1% insulin-transferrin-selenium) at 37°C and 5% CO2. K562 cells were cultured in RPMI 1640 with L-glutamate and 10% fetal bovine serum. HEK293 and HeLa cells were cultured in DMEM high glucose with 10% fetal bovine serum. MCF7 cells were cultured in DMEM high glucose (Hyclone SH30022.01) supplemented with 10% fetal bovine serum (Corning 35-010-CV), 1% penicillin-streptomycin (Hyclone SV30010), and 2 mM L-glutamine (Hyclone SH30034.02).

**Luciferase reporter assay**

H295R cells at 25,000 cells per well (96-well plate) were transfected with 25 ng of dual-luciferase plasmid containing the BoxB or MBE 3' UTR and 25 ng of MSI2 construct using Lipofectamine 2000 (Thermo Scientific) according to the manufacturer's instructions, in parallel with wells for RT-qPCR. Media was refreshed after 24 h. After a further 24 h, cells were harvested and luciferase activity measured using the Nano-Glo Dual-Luciferase kit (Promega) on a GloMax Navigator (Promega) with dual injection.

**Microscopy**

Cells were seeded into 12-well plates (CytoOne) containing 18 mm poly-D-lysine coated German glass coverslips (Fisher Scientific 501929579) and allowed to adhere overnight. Cells were either left untreated or transfected as indicated, washed once with 1X PBS for 5 min at room temperature, and fixed in neutral buffered formalin for 10 min at room temperature. Following fixation, cells were washed three times with PBS for 5 min each, then blocked and permeabilized in CAS-Block containing 0.2% Triton X-100 (Invitrogen 008120) for 5 to 30 min at room temperature with gentle rocking. Primary antibody against MSI2 (Abcam ab76148) was diluted in CAS-T and incubated at 4°C overnight. Samples were washed twice with PBS containing 0.1% Tween-20 (PBS-T) for 5 min each. Secondary antibody (anti-rabbit IgG Fab2 AlexaFluor 647, Cell Signaling 4414) was diluted 1:1000 in PBS-T and incubated for 1 h at room temperature in the dark, followed by two PBS-T washes. Coverslips were mounted onto glass slides (Corning 2948-75X25) using Vectashield mounting medium containing DAPI (Vector Laboratories H-1800-2). Images were captured on a DeltaVision Ultra with a 60X objective in 1.5000 immersion oil over 20 focal planes of 0.4 μm. All images were acquired with identical laser intensity, focal plane number, and step size. Images were deconvolved and quick projected using SoftWorx (GE Healthcare Life Sciences, version 7.2.2) with maximum intensity projection. Image analysis was performed in ImageJ2 (version 2.14.0/1.54p).

**RT-qPCR of reporter mRNA**

H295R cells at 250,000 cells per well (24-well plate) were transfected with 200 ng of dual-luciferase plasmid and 200 ng of MSI2 expression plasmid using Lipofectamine 2000 (Thermo Scientific) according to the manufacturer's instructions, in parallel with wells for the reporter assay. Media was changed after 24 h. After a further 24 h, cells in each well were washed with 500 μL PBS followed by the addition ofand 200 μL RIPA buffer containing with fresh protease inhibitors. was added to each well. Cells were held on ice for 5 min, triturated, and 100 μL of sample was added to 300 μL Trizol LS. RNA was extracted with the microPrep Directzol RNA microprep extraction kit (Zymo Research) according to the manufacturer's instructions. Plasmid DNA depletion was performed as described (Green and Sambrook 2019). Briefly, 3 μL DNase I reaction buffer and 2 μL DNase I were added to 30 μL eluted RNA and incubated for 30 min at 37°C; 2.5 μL of 25 mM EDTA was added and samples incubated for 10 min at 65°C. 100 ng of totaldepleted RNA was used in the iScript cDNA Synthesis Kit (BioRad) according to the manufacturer's instructions, followed by. RT-qPCR withwas performed with iTaq Universal SYBR Green Supermix (Bio-Rad) on a Bio-Rad OPUS 384 instrument. Cycling conditions were 30 s at 95°C, then 40 cycles of 95°C for 5 s and 60°C for 30 s, with signal acquisition after each cycle.

**RT-qPCR of endogenous MSI2 isoform composition**

Total RNA was isolated from HEK293 and H295R cells (Zymo Direct-Zol RNA microprep kit)[kit] and reverse transcribed with a mixture of random hexamers and oligo(dT) and Induro Reverse Transcriptase (NEB) following manufacturer’s instructions [kit], so that transcripts are sampled independently of distance from the poly(A) site. Four primer pairs were used. Pair S1 amplifies only transcripts that contain the 18 amino acid insertion, using a forward primer within the inserted 54 nucleotides. Pair S2 spans the exon 12 acceptor and yields two products, 255 bp with the insertion and 199 bp without it, so a single reaction reports both species. Pairs S3 and S5 share a reverse primer in exon 2 and differ in their forward primer, which lies in the alternative first exon of ENST00000416426 (S3) or in the canonical first exon (S5). GAPDH and RPS20 were amplified as normalizing controls; one was used for normalization and the other to confirm that the normalization was consistent across replicates and cell lines. Reactions were run in 3[n] technical replicates on the Bio-Rad Opus[instrument] as described above.with [cycling conditions], and pProducts were verified by agarose gel electrophoresis.

Isoform composition was calculated from Cq values as 2^-ΔCq ratios between primer pairs that share a denominator. The insertion-containing fraction of full-length MSI2 mRNA was calculated as the S1 signal divided by the summed S2 signal. The alternative first exon fraction was calculated as the S3 signal divided by the sum of the S3 and S5 signals, which is valid because the two pairs share a reverse primer and therefore amplify a common product length from a common target region. Where the S2 reaction was quantified by densitometry, band intensities were divided by product length before taking ratios, because signal from an intercalating dye scales with mass rather than molarity. Lanes of the S2 reaction contained a third product of 279 bp whose abundance tracked the expected heteroduplex frequency, 2p(1-p), across replicates in which the insertion fraction p varied from 0.31 to 0.42; this product was therefore treated as a heteroduplex of the two amplicons and its signal divided evenly between them.

**Western blot**

Cells were induced for 24 h with 2.5 µg/mL doxycycline and collected in 1X Laemmli buffer containing 5% β-mercaptoethanol. Samples were heated for 5 min at 95°C and loaded onto a 15-well, 4%-12% gradient Bis-Tris, 1.0 to 1.5 mm Novex miniprotein gel (Invitrogen), then transferred onto an iBlot mini NC stack nitrocellulose membrane (Invitrogen) using the iBlot2 with the P0 program. Membranes were blocked with 5% nonfat milk in 1X Tris-buffered saline with 0.1% Tween-20 (TBST) for 30 min to 1 h at room temperature, then incubated sequentially with primary antibodies (all 1:2000 in 5% milk TBST) and HRP-labeled secondary antibodies (1:10,000 in 5% milk TBST). Primary antibodies were MSI2 (Abcam ab76148), GFP (Abcam ab290), H3 (Abcam ab18521), and MSI1 (Abcam ab52865). Signals were detected by chemiluminescence on an Azure Biosystems Sapphire biomolecular imager.

**Co-immunoprecipitation**

H295R cells with a stable MSI2 shRNA background containing 3XFLAG-MSI constructs within a doxycycline-inducible loxP locus were plated on 10 cm plates in complete media, grown to 60% confluence, and induced with 2.5 µg/mL doxycycline for 24 h. Cells were collected as described (Wessels et al. 2016). Briefly, media was removed and cells washed with ice-cold 1X PBS, then scraped with a cell lifter into ice-cold 1X PBS and transferred into a pre-weighed 15 mL conical tube. Tubes were centrifuged at 400 × g for 5 min, PBS removed, and the pellet weighed. One volume (w/v) of freshly made PLB buffer was added and cells lysed on ice for 10 min. Whole cell lysate was snap-frozen and stored at -80°C.

The IP was performed as described (Wessels et al. 2016) with modifications. Whole cell lysates were clarified at 20,000 × g for 15 min. The supernatant was transferred to a new tube and total protein quantified with the Pierce BCA 660 nm kit. 320 µg of total protein was used as normalized input per IP. A 5% sample was removed as input for IP-western, and the remainder used as input with pre-washed Sigma Magnetic M2 beads (Sigma M8823; 20 µL slurry or 10 µL packed gel volume per IP) in NT2 buffer. Samples were rotated end-over-end for 3 h at 4°C. Beads were separated using a magnet and the supernatant collected for IP-western. Beads were washed four times with 900 µL NT2 buffer containing 1.5 U RNase I per mg protein input (0.48 U RNase I per 900 µL wash) and resuspended in 50 µL NT2 buffer. Final IP samples were prepared for IP-western by adding 4X Laemmli sample buffer containing β-mercaptoethanol. IP-western was performed as above using mouse anti-FLAG M2 (Sigma F1804) to detect FLAG-tagged constructs, anti-PABP (Abcam ab21060) to detect PABPC1, and rabbit anti-H3 (Abcam ab18521) as loading control.

Similar experiments were performed including endogenous PABP IPs (using ab21060 conjugated to Dynabeads protein A) in H295R cells, and co-overexpression of FLAG-MSI constructs with HA-PABPC1 in HEK293T cells followed by FLAG or HA (Cell Signaling 3724S) IP.

**IP-MS**

H295R cells containing 3XFLAG-MSI2-328, 3XFLAG-MSI2-324, 3XFLAG-MSI2 ΔCLR, or untagged EGFP constructs within a doxycycline-inducible loxP locus were collected as described above. For each sample, three replicates were performed, each containing three 15 cm plates of cells.

The IP was performed as described above. After post-IP washes with RNase I, samples were eluted from the Sigma FLAG M2 magnetic beads with five packed bead volumes (50 µL) of elution buffer (0.1 M glycine, pH 3.5) for 20 min at room temperature with gentle shaking at 500 rpm. The elution was neutralized by addition of 10 µL 0.5 M Tris, 1.5 M NaCl, pH 8.0. An aliquot (2.5%) was saved for IP-western, and the remainder snap frozen in liquid nitrogen and sent to the Fred Hutchinson Cancer Center Proteomics and Metabolomics Core Facility (RRID:SCR_022618) for mass spectrometry. IP-western was performed as above using mouse anti-FLAG M2 (Sigma F1804).

**CLIP-seq**

H295R cells were grown on 15 cm plates in complete media to 60% confluence, washed in 1X PBS, and treated with 300 mJ UV-254 nm (Vari-X-Link) before scraping and pelleting. Three 15 cm plates of cells were used per CLIP-seq replicate.

CLIP-seq was performed as described (Ule et al. 2021) with modifications. Cell pellets were lysed in iCLIP lysis buffer, quantified by BCA, then treated with RNase I at 1.5 and 1.0 U per mg protein lysate input. Dynabeads Protein A (Thermo Fisher) were conjugated to MSI2 antibody (Abcam ab76148) and used for immunoprecipitation of endogenous MSI2 after light RNase I digestion. A 5% size-matched input was taken from the lysate, processed with SP3 beads, and treated in parallel with the CLIP samples following IP. After IP, beads were washed and treated with T4 polynucleotide kinase. Pre-adenylated adapter (/5Phos/AGATCGGAAGAGCACACGTCTG/3Cy55Sp/) was ligated to the 3' ends of RNP complexes with T4 RNA ligase I (NEB M0437M), followed by free adapter removal with 5' deadenylase (NEB M0331S) and RecJ endonuclease (Biosearch Technologies RJ411250NEB M0264S). After eluting the 3'-ligated RNP complexes into 1X Laemmli buffer, samples were run on a 4%-12% Bis-Tris NuPAGE mini-gel in 1X MOPS SDS running buffer for 50 min at 180 V. The gel was transferred to a nitrocellulose membrane using the iBlot2 P0 program, then imaged on the Azure Sapphire biomolecular imager for Cy5 signal. Fluorescent RNP complexes from 55 to 100 kDa were excised, treated with Proteinase K, and RNA extracted. Isolated RNA was used as input into the QIAGEN miRNA-seq library prep kit following the manufacturer's instructions, substituting Superscript IV reverse transcriptase (Thermo Fisher) during cDNA synthesis. Libraries were sequenced on a NovaSeq 6000 for 30 million paired-end reads (2 × 150 bp) each at the University of Colorado Anschutz Medical Campus Cancer Center Genomics Shared Resource Core Facility (RRID:SCR_021984).

**CLIP-seq analysis**

Raw read1 fastq from each sample data was used in the analysis. First UMI were extracted using UMI-tools (PMID: 28100584) (extract --extract-method=regex –bc-pattern=".+(?P<discard_1>AACTGTAGGCACCATCAAT){s<=2}(?P<umi_1>.{12})(?P<discard_2>.*$)") and low quality reads were filtered with FASTX-Toolkit version 0.0.14 (http://hannonlab.cshl.edu/fastx_toolkit/) (fastq_quality_filter -q 10 -p 95). Then adapters were trimmed using cutadapt version 3.5 (DOI:10.14806/ej.17.1.200) (-g CTACACGACGCTCTTCCGATCT -j 0 -q 15,10 -m 20) and reads were collapsed based on UMI using custom python and perl scripts. Then repetitive elements were removed and remaining reads were aligned to the reference genome using STAR version 2.7.9a (PMID: 23104886) (--outSAMunmapped Within --outFilterMultimapNmax 1 --outFilterMultimapScoreRange 1 --outSAMattributes All --outSAMtype BAM Unsorted --outFilterType BySJout --outFilterMismatchNoverReadLmax 0.04 --outSAMattrRGline ID:foo --alignEndsType EndToEnd --outSAMmode Full). For endogenous MSI2, peaks were called using omniCLIP version 0.20 (PMID: 30384847) and annotated using custom R scripts.

**Transcript-region distribution of CLIP reads.** Reads from MSI2-324 and MSI2-328 CLIP libraries were assigned to transcript regions using their aligned blocks. Each read contributed a total weight of one, divided equally among its distinct overlapping annotation categories. Category weights were summed and divided by the total number of eligible aligned reads in each library. Fractions were calculated separately for two biological replicates per isoform and summarized by their arithmetic mean, without size-matched input normalization. Intergenic and other annotations were combined within each replicate where indicated. The intron/3′ UTR-only plot retained the original denominator without renormalization.

**Normalized intronic peak distributions.** MSI2 peak summits were assigned to overlapping introns on the same strand, selecting the longest matching intron when multiple assignments were possible. Introns longer than 500 nt were retained, yielding 20,364 summits in 9,565 introns. Summit positions were oriented from the intron’s 5′ to 3′ end, normalized by intron length, and grouped into 100 equal-width bins. Each represented intron contributed equal total weight, divided among its assigned summits. Null profiles were generated from 500 randomizations that independently repositioned summits within their assigned introns while preserving intron lengths and summit counts and excluding 12.5 nt at each end. Observed profiles were plotted without smoothing alongside the null median and pointwise 10th–90th and 25th–75th percentile bands.

**Splice junction analysis**

Split reads spanning the shared MSI2 exon 11 donor (chr17:57652162, hg38) were counted for the two annotated 3' acceptors, at chr17:57674917 (generating the 18 amino acid insertion) and chr17:57674971 (not generating it). Acceptor usage was expressed as the fraction of reads at the two acceptors that use the insertion-generating acceptor. GTEx v11 and TCGA values were computed from STAR junction counts, restricting to samples with at least 20 reads spanning the donor (10,765 of 19,788 GTEx samples). Developmental and cell-type analyses used the Snaptron srav3h compilation of public human RNA-seq, restricting to samples with at least 50 spanning reads; samples were classified from SRA run and sample metadata, and per-study values were pooled across samples by summing reads rather than averaging per-sample fractions. Values from the two compilations are compared directly because they agree closely on the same samples: across GTEx, acceptor usage was 30.9% from STAR junction counts and 29.2% from the Snaptron gtexv2 compilation (medians, samples with at least 20 spanning reads). Category-level values are medians across samples, with 95% confidence intervals obtained by bootstrap resampling of samples (10,000 resamples). Paired comparisons across studies used the Wilcoxon signed-rank test.

**Tumor and normal comparisons**

TCGA tumor samples and matched normal samples were analyzed with the same junction-counting procedure. Cohorts were included if at least five matched normal samples passed the read threshold, which retained 13 cohorts. Cohort-level comparisons used the Mann-Whitney test without correction for multiple comparisons, because the cohorts represent independent tumor types rather than repeated tests of one hypothesis. The paired analysis was restricted to the 291 patients from whom both a tumor and a normal sample were available and used the Wilcoxon signed-rank test. For the survival analysis, acceptor usage was converted to a z-score within each cohort and the four cohorts with significantly elevated usage were pooled; overall survival was censored at 10 years and the hazard ratio estimated by Cox proportional hazards regression with Breslow handling of ties.
